# Stealth fluorescence labeling for live microscopy imaging of mRNA delivery

**DOI:** 10.1101/2020.07.01.172767

**Authors:** Tom Baladi, Jesper R. Nilsson, Audrey Gallud, Emanuele Celauro, Cécile Gasse, Fabienne Levi-Acobas, Ivo Sarac, Marcel Hollenstein, Anders Dahlén, Elin K. Esbjörner, L. Marcus Wilhelmsson

## Abstract

Methods for tracking of RNA molecules inside living cells are critical to probe their dynamics and biological functions, but also to monitor delivery of therapeutic RNA. We here describe a method for fluorescence labeling of RNAs of any length, via the enzymatic incorporation of the minimally perturbing and intrinsically fluorescent tricyclic cytosine analogue tC^O^. Using this approach, we demonstrate incorporation of tC^O^ in up to 100% of all natural cytosine positions of a 1.2 kb mRNA encoding for the histone H2B fused to GFP (H2B:GFP). The resulting transcript is fully compatible with both in vitro transcription and subsequent in cell translation. Spectroscopic characterization of the in vitro transcribed mRNA, shows that the incorporation rate of tC^O^ is on par with cytosine, facilitating efficient labeling and controlled tuning of labeling ratios for different applications. Using live cell confocal microscopy and flow cytometry, we show that the tC^O^-labeled mRNA is efficiently and correctly translated into H2B:GFP upon electroporation as well as lipid-mediated transfection of human Huh-7 cells; correct translation was further confirmed in cell-free systems. Importantly, the spectral properties of the tC^O^-modified transcripts and their translation product, in this case H2B:GFP, allow for their straightforward and simultaneous visualization in live cells.

---

RNA is a key molecule of life and a main active player of the central dogma of molecular biology. RNA is a crucial regulator of gene expression via for instance micro (mi)- and small interfering (si)-RNA and through its intrinsic catalytic activity, RNA plays a fundamental role in biology. It has, for these reasons, also emerged as a highly promising and versatile drug modality; RNA therapeutics have the potential to modify cellular function at the translational level, which opens up entirely new opportunities to address previously undruggable targets.^1^ An increased molecular and mechanistic knowledge of the biological processes involving RNA is therefore vital for understanding diseases and to treat them. A growing body of evidence suggests that the key to unleashing the full potential of RNA-based drugs lies in understanding the processes and mechanisms of cell uptake and endosomal release.^2^ Regardless of the endocytosis mechanism,^3^ the delivery of a nucleic acid cargo to the cytoplasm always relies on endosomal escape, the understanding of which, despite extensive investigations, remains elusive^4,5^ much due to the challenges associated with visualizing RNA released to the cytosol. In this context, tracking of endogenous and exogenous (therapeutic) RNAs inside cells, including their translocation, localization, splicing and degradation, is of great importance. Recent advances have resulted in the development of a broad spectrum of tools and probes by which RNA can be analyzed and quantified, but they generally involve heavily modified oligonucleotides with properties that are significantly different from natural ones, potentially resulting in loss of ability to be correctly recognized and processed by the enzymatic machineries of cells. Hence, the development of methods for universal, minimally perturbing and internal RNA labeling schemes that are compatible with live cell imaging and fluorescence microscopy, may become as crucial for the RNA field as the discovery and understanding of the Green Fluorescent Protein (GFP) has been for proteins.^6–8^

A drawback of existing fluorescence-based technologies for studying cellular localization of RNA is that they primarily rely on highly amphiphilic and/or bulky external fluorescent constructs which could impair motility and perturb localization of the RNA^9^ and its molecular interactions with proteins and membrane constituents.^10^ In addition, a majority of these technologies are incompatible with live cell imaging;^11^ Fluorescence In Situ Hybridization (FISH), for example, can be used to visualize and quantify ribosome-mRNA interactions with single mRNA transcript resolution,^12^ but it is limited to fixed cells. Förster Resonance Energy Transfer (FRET) is another highly attractive option for obtaining information on interactions, localization and relative amount of mRNA strands. It has for instance been shown that antisense oligonucleotides (ASOs) functionalized with FRET-pairs that light up only upon interaction with a specific sequence can be used to probe for specific mRNA targets. However, binding of multiple FRET-pair-labeled ASOs to an RNA may perturb its native function by sterically blocking protein binding sites and translation; there is also a risk that the RNA target sequence is inaccessible due to complex tertiary structures of the mRNA or bound RNA proteins, causing the light-up ASO probes to report false negatives. In addition, although this approach is in principle compatible with live cell imaging, it has so far mostly been used in fixed cells, combined with FISH, to provide highly detailed, yet only static, information.^13,14^ In the few examples where RNA dynamics has been visualized in live cells, the probes were typically covalently conjugated to a vehicle, such as a DNA nanocages,^15^ or gold nanoparticle,^16^ before delivery to cells. Another emerging strategy for labeling RNAs is via fusion to aptamer sequences that will bind and activate fluorescent dyes in situ,^17^ but it was shown that the site of insertion of such constructs onto the RNA of interest can influence the translation efficiency and protein interactions of the latter.^18^ To reduce the risk of interfering with enzymatic processes, chimeric mRNAs have also been used, carrying multiple stem-loop structures in the 3’-UTR and downstream of the STOP codon, which serve as binding sites for fluorescent fusion proteins.^19,20,21^ Whilst versatile, all these labeling strategies impact profoundly on the RNA molecular weight and there is hence a risk of perturbed motility and intracellular transport. Additional less perturbing strategies for labeling mRNA outside of the coding region include azido-functionalized 5′-cap analogues that can be incorporated using in vitro transcription and further “clicked” to a dye inside live cells^22^ and 3’-polyA tail tags based on azido-functionalized adenosine triphosphate analogues.^23^ Click-chemistry approaches have also been used to introduce fluorophores into the coding sequence of an mRNA post-transcription.^24^ Despite progress in these areas, external, molecular dyes, such as the cyanines Cy3 and Cy5 remain the most common labeling choice.^25,26^ Design and implementation of smaller and uncharged fluorophores for intrinsic native-like labeling of RNAs, compatible with both transcriptional and translational machineries, will therefore represent a major step forward. A first step in this direction has been taken by Ziemniak et al., who developed fluorescent 5′-cap analogues compatible with both transcription and translation.^27^ However, the resulting RNA products have not been visualized inside cells, limiting the applicability of the method.

Fluorescent nucleobase analogues (FBAs) have emerged as attractive labels for both DNA and RNA and we have led the development towards increased brightness values and tuning of wavelengths of excitation and emission to facilitate their utilization in fluorescence microscopy, something that until now has not been accomplished.^28–30^ Since they are internal fluorophores (i.e. involving a modification which primarily affects the oligonucleotide’s interior), they are less functionally perturbing than other probes. Furthermore, their design enables them to retain the normal base-pairing and -stacking of the target nucleic acid. These native-like fluorescent labels thereby open new possibilities, not only to track the nucleic acid of interest but also to use their fluorescence read-outs to obtain detailed information regarding nucleic acid structure and behavior. We have, for example, designed and implemented the first FBA interbase FRET-pair both in DNA^31^ and RNA,^32,33^ enabling detailed distance and orientation monitoring over more than one turn of the duplex. Fluorescent RNA base analogues have also been used to study the rate of codon:anticodon base-pair formation during translation^34^ and a few studies demonstrate their use as substrates for in vitro transcription reactions, demonstrating T7 RNA polymerase mediated incorporation the fluorescent tricyclic cytosine analogue tCTP into *ca.* 800 nucleotide (nt) non-coding RNA strands^35^ or the production of short transcripts incorporating fluorescent isomorphic guanine^36,37^ and uridine^38,39^ analogues, respectively. Yet again, none of these studies have explored if it is feasible to probe the labeled RNAs by their fluorescence in a live cell context.

In this study, we have developed a method to incorporate fluorescent base analogue internal labels at desired ratios into a 1.2 kb long coding mRNA and demonstrated the functionality of the resulting fluorescent transcript both in vitro and by live cell confocal microscopy imaging in a drug delivery context. Our method, that presumably is applicable to most mRNAs, allows for efficient in vitro transcription and, remarkably, an effective in-cell translation of full-length mRNAs internally labeled with the fluorescent tricyclic cytosine tC^O^ at all cytosine positions. The absorption of tC^O^ is centered at 369 nm (*ε*_369_ = 9370 M^−1^ cm^−1^) and it fluoresces intensely in single-stranded (ss) RNA (<*Φ*_*F*_> = 0.24), with maximum emission at 457 nm.^32^ We demonstrate that the tC^O^-mRNA is readily imageable inside live cells using the standard 405 nm laser line of a confocal microscope for excitation and, moreover, that we can simultaneously follow the emergence of the transcript’s protein product – histone protein H2B fused to GFP (H2B:GFP), which localizes to the nucleus (Figure 1). We also present an alternative enzymatic labeling strategy for RNAs of any length in which we demonstrate the conjugation of tC^O^ to the end of RNA sequences. Since the tC^O^ label is compatible with biological processes that RNA participates in, it holds a great potential to become a powerful imaging tool in live cell microscopy. Specifically, we envision this approach to facilitate drug delivery studies, enabling effective visualization of cellular uptake, endosomal release, exosomal loading and trafficking of mRNA.

**Figure 1.**
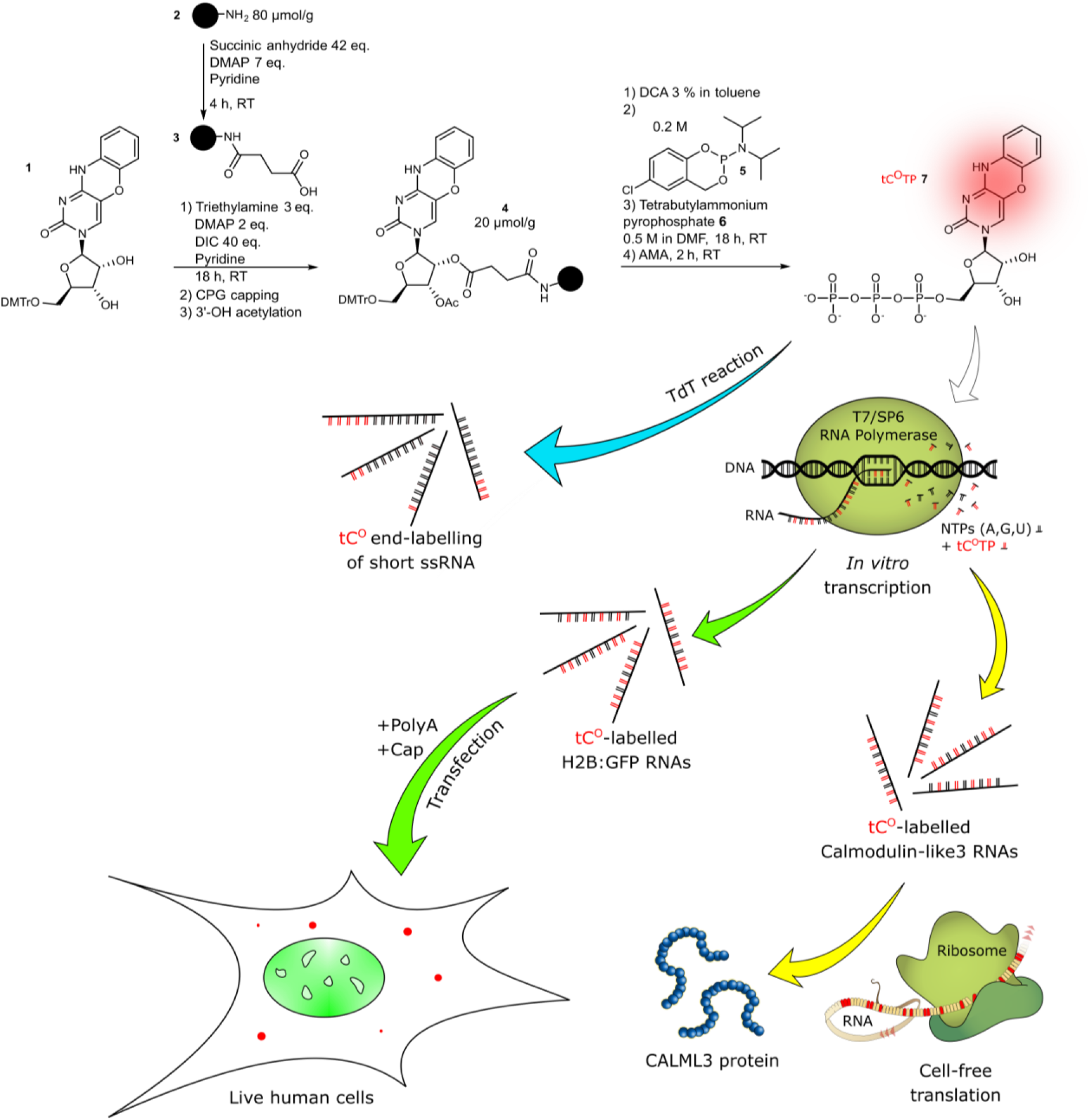
Schematic outline of the study. **Blue pathway**: End-labeling of ssRNA sequences using a terminal deoxynucleotidyl transferase (TdT). **Green pathway**: Cell-free transcription into H2B:GFP mRNA, transfection into human cells and live-cell imaging of both the GFP protein (translation product) and tC^O^ (mRNA). **Yellow pathway**: Cell-free transcription into Calmodulin-like3 RNA followed by cell-free translation into the CALML3 protein. Color code: tC^O^ in red, GFP in green, native nucleosides in black or beige.

## Results and Discussion

### Synthesis of the tricyclic cytosine analog (tC^O^) triphosphate

Since the Yoshikawa^40^ and Ludwig-Eckstein^41^ conditions were published in 1969 and 1989, respectively, a plethora of alternatives have been developed for the triphosphorylation of nucleosides.^42–45^ The abundance of literature on this matter well illustrates the “hit-and-miss” aspect of triphosphate synthesis. Both the triphosphorylation process and the laborious purification of the very polar final products are liable for the typically low synthesis yields.

To improve the reaction yield and facilitate purification of the final product during the synthesis of modified RNA triphosphates, we opted for a synthetic scheme (Fig. 1) starting from a solid-supported ribonucleoside; this approach has surprisingly only been reported for unmodified nucleobases with 2′-OMe protection.^46,47^

The final reaction, relying on well-established phosphoramidite chemistry, involves the use of *Cyclo*Sal-phosphoramidite **5** and bis(tetrabutylammonium) dihydrogen pyrophosphate **6** (Fig. 1), which are prepared from commercially available 5-chlorosalicylic acid and pyrophosphoric acid. This method was developed by Meier et al.^48^ to achieve efficient 5′-triphosphorylation of short solid support-bound DNA and RNA oligonucleotides in moderate to good yields. Interestingly, it was originally used for solution-based synthesis of unmodified ribonucleosides and deoxyribonucleosides,^49^ but has progressively been combined with solid supports to facilitate purification of the products.^50^ In our hands, both Krupp’s^46^ and Meier’s^48^ solid-phase triphosphorylation methods yielded the product, though Meier’s gave a higher yield. To the best of our knowledge, the *Cyclo*Sal-phosphoramidite method on solid support described herein has never been applied to a single ribonucleoside before, let alone to produce a modified base analogue.

Briefly, long-chain alkylamino controlled-porosity glass (CPG) support was functionalized with a succinyl moiety. Protected ribonucleoside **1** was synthesized according to a method by Füchtbauer et al.^32^ and attached to the succinylated support via ester bond formation (Fig. 1). The resulting ribonucleoside **4** on solid support could be stored in the dark at room temperature, with no degradation observed over three months. Interestingly, intermediate **4** could also be synthesized via succinylation of ribonucleoside **1** in solution followed by coupling with the amino support **2**. Subsequently, triphosphorylation of ribonucleoside **4** was performed using the *Cyclo*Sal method. After triphosphorylation, support-bound triphosphate was deacetylated and cleaved from the CPG support using ammonium hydroxide/methylamine (AMA) for two hours at room temperature. Subsequent reverse phase or ion-exchange chromatography allowed the desired triphosphate **7** in a triphosphorylation yield of 60% and with a high UV purity of 99%. Importantly, up to 85% of the unreacted nucleoside **1** could conveniently be recovered by precipitation from the first reaction crude, compensating for the low loading achieved. Considering this, our triphosphorylation method gives an overall yield of up to 30%, which is higher than most solution-based alternatives.

### Cell-free enzymatic incorporation of tC^O^ into RNA

The tC^O^ triphosphate (tC^O^TP) was used to produce fluorescently labeled RNA via two different enzymatic pathways. First, we performed tail-labeling of short RNA oligonucleotides using the terminal deoxynucleotidyl transferase enzyme (TdT, Fig. 1, blue arrow). This template-independent DNA polymerase appends both natural and modified nucleotides at the 3’-end of single-stranded oligonucleotides but, unlike most DNA polymerases, TdT displays a characteristic low discrimination for deoxyribo-over ribonucleoside triphosphates, enabling modification also of RNA. ^51,52^ The TdT-mediated tail-labeling of a 17-mer ssRNA (TdT1, see Supplementary Table 1 for sequence) resulted in successful addition of multiple tC^O^s (from one to more than 25) and with near full conversion of the RNA primer (Fig. 2b). The TdT usually displays no strict metal cation preference,^51^ which was confirmed also in our case, with equally effective addition of tC^O^s with Co(II), Mg(II), or Mn(II) as enzymatic co-factor. Thus, even though RNA nucleotides have been reported to be relatively poor substrates for TdT,^53,54^ tC^O^TP appears to be well tolerated by this enzyme. We next demonstrated that it is possible to use this approach to tail-label also longer RNA sequences, enzymatically adding tC^O^s to a 50 nt ssRNA (TdT2, Supplementary Table 1). The labeling was effective also in this case resulting in near full conversion of the primer, but led to a significantly higher degree of product uniformity; the main products contained one or two tC^O^ incorporations (Supplementary Fig. 1). The fluorescence originating from a tC^O^ tail-labeled oligonucleotide was clearly observable by naked eye upon UV irradiation (Fig. 2c). This result shows that the tricyclic tC^O^ analogue can be effectively read by the TdT enzyme and thereby incorporated into RNA of any length providing site-specific fluorescence end-labeling. This opens up for advantageous possibilities for dual labeling approaches, where combinations of tC^O^ with other base- or backbone-modified fluorescent markers could be used to monitor for example in vivo stability.

**Figure 2.**
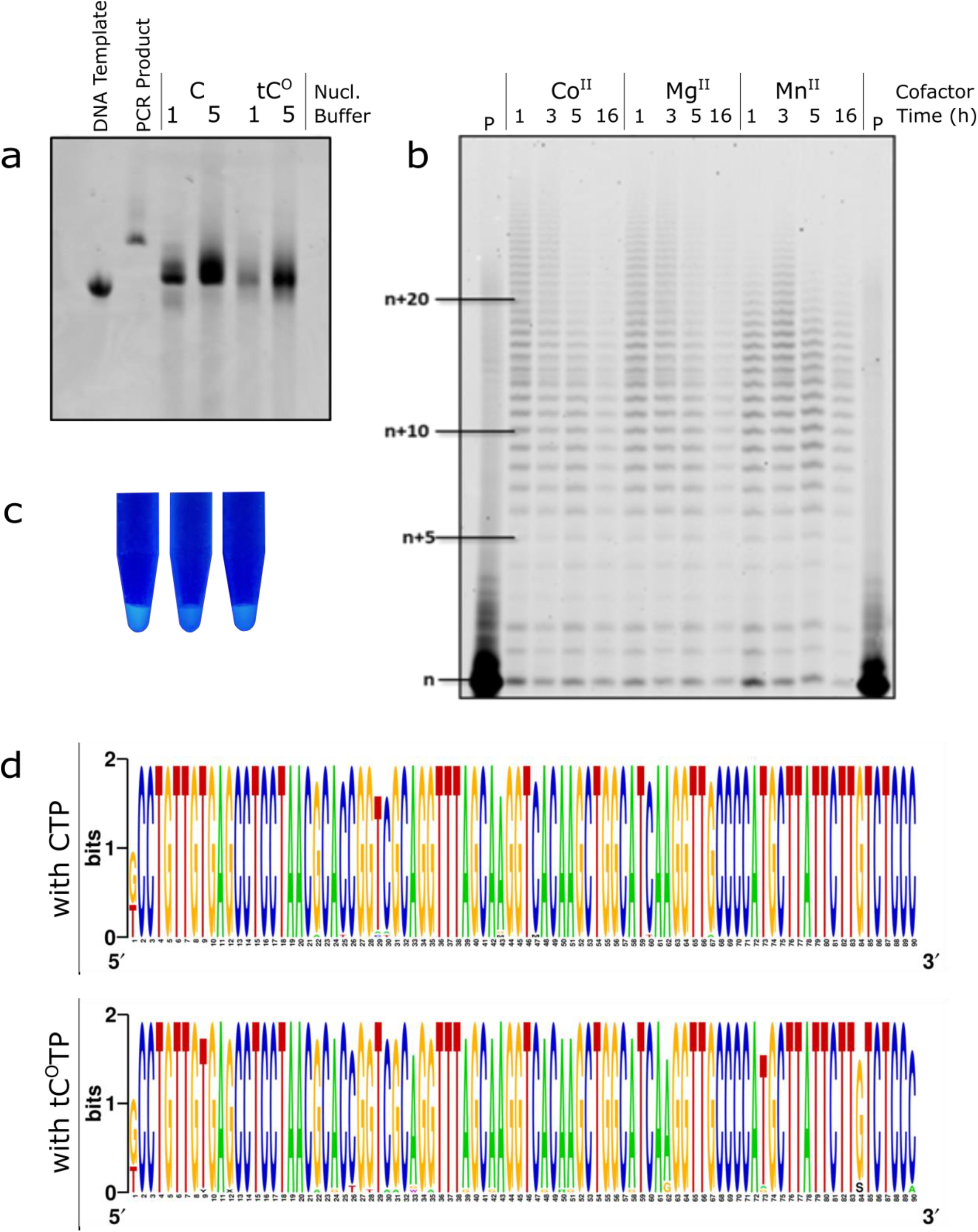
Enzymatic incorporation of the modified triphosphate by in vitro transcription on short templates, sequencing of reverse transcriptase products and TdT-mediated labeling. a) Gel image (PAGE 10%; visualization by midori green) of RNA products from 6 hours of T7 RNA polymerase assisted transcription using the D1 Library as DNA template. See Methods for buffers composition. b) Gel image (PAGE 20%; visualization by phosphorimager) of the products from TdT-mediated end-labeling of oligonucleotide TdT1 using tC^O^TP. The lowest band labeled “n” represents the unreacted TdT1 oligonucleotide and all above bands are different products with increasing additions of tC^O^ to the tail. P: control reaction in absence of polymerase. c) Picture of UV-light irradiated solutions of purified TdT3 RNA end-labeled with tC^O^. d) Sequence logo from the cloning-sequencing protocol of reverse transcription products from the modified T12.3 RNA (top: products from a 6 h transcription reaction with CTP; bottom: products from a 6 h transcription reaction with tC^O^TP).

Next, we explored the capacity of T7 RNA polymerase to incorporate tC^O^TP into short (20 nt) RNA transcripts. Under optimized buffer conditions, we observed successful transcription upon complete replacement of canonical CTP with tC^O^TP, yielding the full-length 20 nt long transcripts without any signs of premature transcription termination (Supplementary Fig. 2a). Longer 90 nt DNA templates (T12.3 and D1 Library, Supplementary Table 1) were also compatible with the T7 RNA polymerase/tC^O^TP system (Fig. 2a and Supplementary Fig. 3), confirming that tC^O^TP is readily accepted by the T7 polymerase as a substrate. Stengel et al. have shown similar results using the structurally related tricyclic cytosine analogue tC, which carries a bulkier sulfur atom instead of an oxygen at position 5 of the pyrimidone ring, but the reaction proceeded with a high 40–300 discrimination factor against the formation of mismatched tC-A base pairs.^35^ We therefore evaluated the propensity of tC^O^TP to be misinserted opposite to template deoxyadenosines using the templates DNA3 and DNA4 (Supplementary Table 1), which contain no guanosine and a single adenosine downstream of the promoter sequence. Transcription reactions were run in the presence of either CTP or tC^O^TP and absence of UTP (Supplementary Fig. 2b). With the DNA4 template, where the mismatch position is located close to the T7 promoter sequence, tC^O^TP in fact induced slightly fewer aborted transcripts and more full-length transcripts were produced than with the native CTP. However, when using the DNA3 template, where the mismatch position is located further down the sequence, no noticeable difference was observed in the reactivity of tC^O^TP and CTP. To more thoroughly evaluate the fidelity of incorporation of the modified nucleotide, we then converted a tC^O^-containing RNA oligonucleotide obtained from in vitro transcription of template T12.3 to unlabeled cDNA using M-MLV reverse transcriptase (Supplementary Fig. 3). After PCR amplification and A-tailing reaction, the resulting amplicons were ligated to the pGEM T vector and transfected into beta 2033 competent E. coli cells. Plasmids stemming from white colonies were then subjected to Illumina sequencing. Analysis of the multiple alignments of the sequences originating from these transcription reactions revealed a two-fold higher frequency of misincorporation when tC^O^TP was used instead of CTP (28 vs. 16 points mutations in respectively 34 and 38 analyzed sequences; Supplementary Fig. 4-5) and only some significant A→G transversions at positions 24, 33, 39 and 62 (Fig. 2d) Altogether, this shows that the fluorescent tC^O^TP nucleotide can be processed by both TdT and T7 RNA polymerases, with the latter only marginally increasing the incorporation error rate in up to 50 nt long RNAs. In this sense, tC^O^TP behaves better or on par with a majority of previously modified nucleotides,^35,36,55-57^ including bulkier triphosphate analogues.^58,59^ It has been reported that the T7 RNA polymerase binds the incoming nucleotide substrate in an open conformation,^60^ which could enable it to accommodate our modified base.

### Cell-free enzymatic incorporation of tC^O^ into mRNA transcript

We next used cell-free in vitro transcription to produce fluorescent full-length messenger RNA (mRNA), from a DNA template encoding for H2B histone protein fused to GFP (H2B:GFP). The template was codon optimized (Supplementary Fig. 6) to limit the number of C repeats, since adjacent tC^O^ moieties in oligonucleotides are known to self-quench, which could cause loss of proportionality between the degree of incorporation and the observed fluorescence intensity^61^ in the transcript characterization (vide infra). We observed efficient transcription and tC^O^ incorporation using two different bacteriophage RNA polymerases, T7 and SP6 (Fig. 1, first green arrow) at tC^O^TP/canonical CTP ratios ranging from 0 to 100% (full replacement), as demonstrated by agarose bleach gel electrophoresis (Fig. 3a for T7 and Supplementary Fig. 7a for SP6). All RNA transcripts appear as one single band on the gels, with a migration corresponding to the expected 1247 nt mRNA product (H2B:GFP), demonstrating that full length mRNA was formed. The tC^O^-containing mRNA bands could be directly visualized upon 302 nm excitation using a conventional gel scanner (Fig. 3a); the increasing band intensities with increasing tC^O^TP/CTP reaction ratio supported successful concentration-dependent incorporation of tC^O^. Re-visualization of the gel after ethidium bromide staining (Fig. 3a, right) revealed similar amounts of RNA in all reactions, providing a qualitative indication of that tC^O^ incorporation does not reduce the in vitro transcription reaction yield. Furthermore, no shorter transcripts were observed, strengthening the observation in Fig. 2d that the T7 polymerase processes tC^O^TP correctly and without premature abortion. Higher order bands are apparent in all lanes of the gel (Fig. 3b and Supplementary Fig. 7b) but were removed upon heat denaturation, suggesting the presence of RNA secondary structures. Notably, this feature appears independently of the CTP/tC^O^TP-ratio, indicating that the effect is not specific to the modified cytosine base. Importantly, we demonstrate that tC^O^ can be successfully incorporated into full-length RNA transcripts even under conditions where all canonical CTP is replaced with tC^O^TP (0% CTP; i.e. 100% C-labeling efficiency).

**Figure 3.**
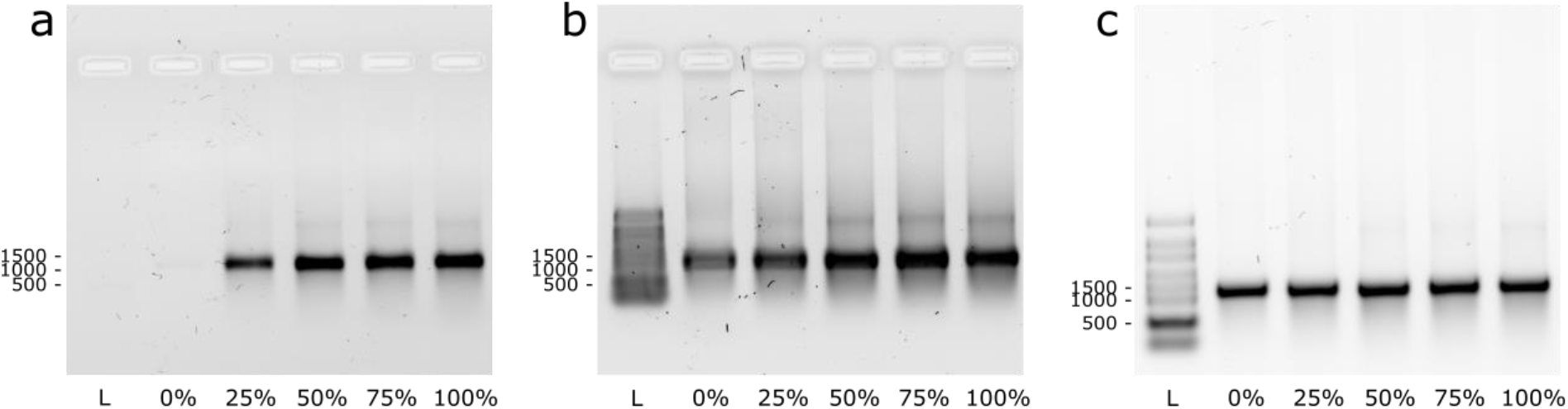
Incorporation of tC^O^ into full length mRNA by T7 RNA polymerase assisted in vitro transcription. Denaturing agarose bleach gels showing RNA transcripts formed at five different tC^O^TP/CTP ratios (0-100%). Direct visualization of tC^O^ fluorescence (a) and after ethidium bromide staining (b). RNA samples were heat-denatured (65 °C for 5 min, 1.5% bleach in the gel) prior to loading. (c) Same RNA transcripts upon harsher denaturation (70 °C for 10 min., 2% bleach in the gel). The RiboRuler High Range RNA ladder was used.

### Spectroscopic characterization of in vitro synthesized tC^O^-modified RNA transcripts

We used a spectroscopic approach to quantify the incorporation efficiency of tC^O^TP, compared to the canonical CTP. To enable this, all RNA transcripts were purified using a Monarch RNA Cleanup kit, ensuring complete removal of unreacted tC^O^TP. Absorption spectra (Fig. 4a) showed the appearance of a band centered at ca. 370 nm in samples with the incorporated cytosine analogue tC^O^, consistent with the spectral profile of this FBA (Fig. 4a).^32^

**Figure 4.**
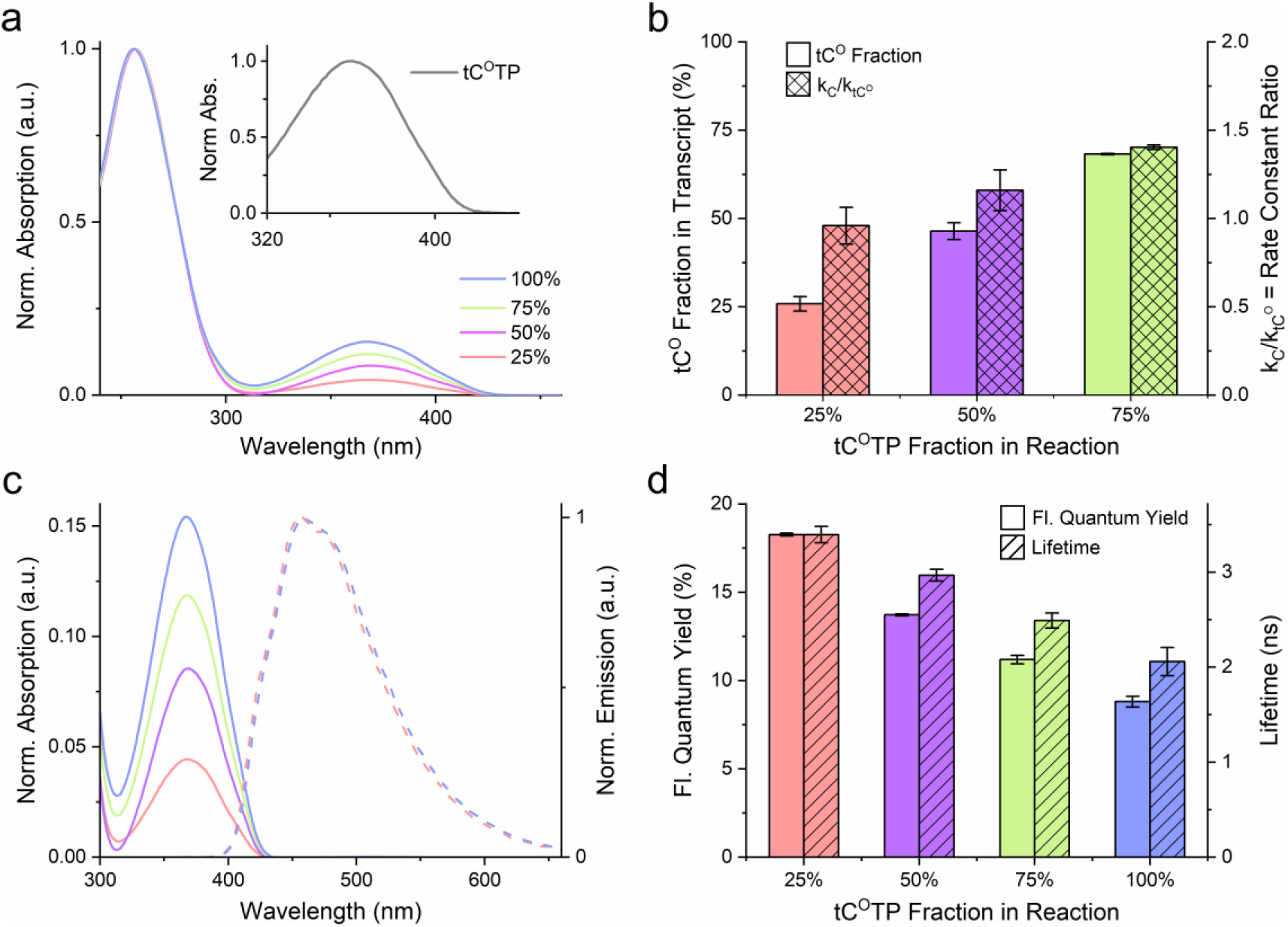
Spectroscopic characterization of in vitro synthesized tC^O^-modified RNA transcripts. Four reactions charged with different molar fractions of tC^O^TP (blue: 100%, green: 75%, magenta: 50%, and red: 25%) in the total cytosine triphosphate pool (tC^O^TP + CTP) were performed. The product transcripts were purified to wash out unreacted triphosphates prior to characterization. All reactions were performed as independent duplicates and the results are presented as mean ± standard deviation. **a)** UV-vis absorption spectra normalized to *A* = 1 at the RNA band, ca. 260 nm. Inset: tC^O^TP absorption normalized to *A* = 1 at the tC^O^-band λ*_max_* (360 nm). **b)** Plain bars: Fraction of incorporated tC^O^ (relative to the total amount of incorporated cytosines, i.e. tC^O^ + C) in the transcripts. Checkered bars: Ratio of first-order reaction rate constants for CTP vs. tC^O^TP consumption. **c)** Solid lines: UV-vis absorption spectra (normalized to *A* = 1 at the RNA band, ca. 260 nm) showing the tC^O^-band centered at 368–369 nm. Dashed lines: Emission spectra normalized to *I* = 1 at *λ_max_* (457 nm and 459 nm for the 25% and 100% transcript, respectively). For clarity, the emission spectra for the 50% and 75% reactions were omitted. **d)** Plain bars: Fluorescence quantum yields. Striped bars: Fluorescence lifetime.

By relating the absorption of the purified RNA transcripts at 260 nm, which reflects their total concentration, to the absorption at 370 nm (emanating exclusively from tC^O^), we could determine the relative rate constants for the incorporation of CTP and tC^O^TP (k_C_ and k_tC^O^_, respectively, see Methods for details). The calculated quotients k_C_/k_tC^O^_ (Fig. 4b) are close to or slightly above unity (0.96–1.4, Fig. 4b), demonstrating that the T7 polymerase displays no substantial preference for the canonical CTP over tC^O^TP. This supports the idea that the tricyclic chemical modification is indeed minimally perturbing in the transcriptional process. Our observation of equal incorporation efficiencies for tC^O^TP and CTP differs from other published studies where the corresponding incorporation ratio was found to be lower than 0.6 (and down to 0.13) for d(tC^O^TP) incorporation into DNA, and analogues d(tCTP) into DNA and tCTP into RNA.^35,62,63^ Consequently, we find that the resulting percentage of tC^O^ in the transcripts overall match very well with the percentage of tC^O^TP added to the transcription reaction (Fig. 4b). This, importantly, demonstrates that the labeling density can be tuned in a flexible and facile way to produce transcripts with labeling ratios tailored to match different applications.

We next examined the emissive behavior of tC^O^ in the mRNA transcripts exploring its relation to the tC^O^TP fraction added to the initial reaction mixture. A substantial decrease in fluorescence quantum yield (from 0.18 to 0.09, Fig. 4d) was observed with increasing tC^O^ incorporation, despite codon optimization to minimize the number of adjacent C positions (vide supra). This was accompanied by a decrease in fluorescence lifetime (from 4.3 ns to 3.2 ns) and a slight red-shift of the emission spectrum (*ca.* 4 nm, Fig. 4c). We ascribe this to electronic coupling of molecular states of the tC^O^ fluorophore and a self-quenching effect caused by the increasing concentration of vicinal tC^O^s (vide supra and Supporting Fig. S7). Self-quenching is an established phenomenon for low Stokes shift fluorophores in general^61^ and a similar observation was made by Stengel et al. in previous DNA polymerization assays using d(tC^O^TP).^62^ Importantly, this quenching effect at high tC^O^ fractions is, more than well, balanced by the large overall number of incorporations and does not prevent visualization of the mRNA, even for transcripts where all C positions are replaced by tC^O^.

### Translation of tC^O^-labeled mRNA in bacterial lysates

In order to verify the functionality of the tC^O^-labeled mRNA transcripts we investigated translation of in vitro transcribed Calmodulin-3 mRNA in cell-free conditions using bacterial lysates (Fig. 1, yellow arrows). The labeled mRNA was produced using a commercial Calmodulin-3 DNA template plasmid using the same tC^O^TP/CTP ratios as for the H2B:GFP encoding mRNA (0 to 100% of tC^O^TP). After RNA purification and cell-free translation, we confirmed the presence of Calmodulin-3 by Coomassie staining (Fig. 5a) as well as Western Blot (Fig. 5b). Satisfactorily, we observed stable protein expression levels when increasing the tC^O^ content of the transcripts, ranging from 80%–137% of the non-labeled control (Figure 5c). This result suggests that tC^O^, unlike other sugar-^57^ or base-modified^64^ nucleotide analogues, does not impair ribosomal processing in cell-free systems.

**Figure 5.**
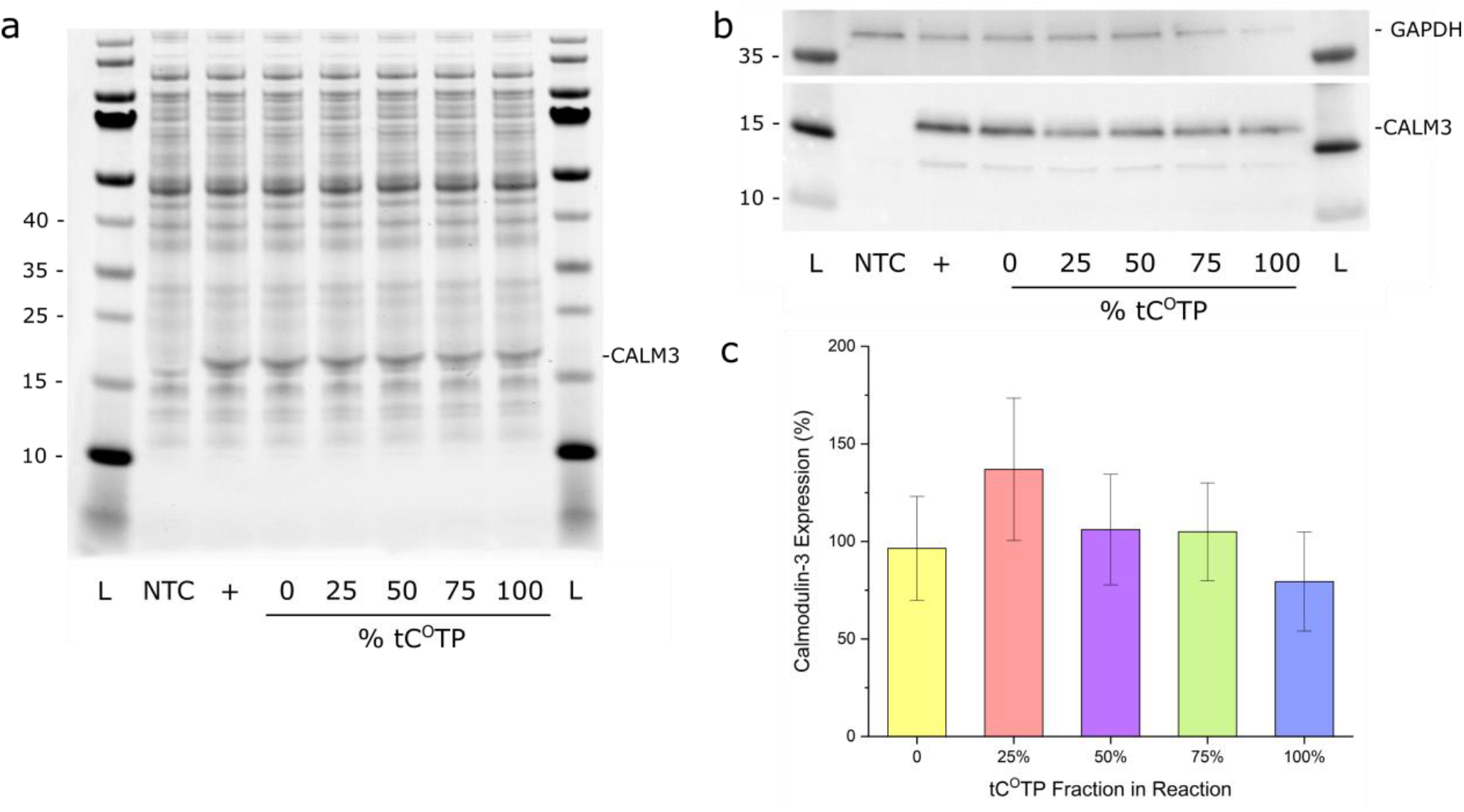
**Cell-free translation.** a) Coomassie staining and b) Western Blot (WB) of the in vitro translation reactions. NTC: no template control; +: kit template RNA control. The PageRuler Prestained Protein Ladder was used. c) Quantification by WB and densitometry analysis (mean % of control ± standard deviation of three replicates).

### Translation efficiency of tC^O^-labeled mRNA in human cells

We used electroporation to introduce the in vitro-transcribed tC^O^-labeled mRNA transcripts into human neuroblastoma SH-SY5Y cells (Figure 1, second green arrow). Taking advantage of the fact that the mRNA encodes for a fluorescent fusion protein with nuclear localization (H2B:GFP), the translation could conveniently be detected via the product’s fluorescence (Fig. 6). To improve stability and reduce cytosolic degradation, the mRNA was capped with a 5′-Cap 0 analogue and 3’-protected by poly-adenylation (by ca. 300 nt). We showed by live cell confocal fluorescence microscopy and flow cytometry analysis that nuclear GFP fluorescence could be detected in 32%, 25%, 18%, and 12% of the cells 24 hours post the electroporation of mRNA containing 25%, 50%, 75% and 100% of tC^O^ (Fig. 5a and Supplementary Fig. 8a). In comparison, the transfection efficiency with unmodified mRNA was 46%. This provides the first observation that an FBA-modified RNA transcript can be accurately and efficiently translated by human ribosomal machineries, resulting in a correctly localized and folded protein product. Using flow cytometry, we quantified the levels of H2B:GFP fluorescence in the cells (Fig. 5b), observing a decrease in mean cellular H2B:GFP fluorescence intensity upon increasing the percentage of tC^O^ in the transcript (approximately one order of magnitude difference between 0 and 100% of tC^O^ (Fig. 5a and 5f)). This suggests that at least under these conditions, mRNA translation, as opposed to transcription, is somewhat impeded by the tC^O^ modification, especially at the highest degree of incorporation. Importantly, we found no evidence of mRNA-induced cell toxicity up to 24 hours post-electroporation (Supplementary Fig. 8g). Of significant note is that the mean fluorescence intensity of translated protein in cell cultures electroporated with a commercial enhanced GFP-encoding mRNA tagged with Cy5 via conjugation to UTPs (ca. 25% of all U positions of this mRNA are labeled) was only 17% of that in cell cultures electroporated with the corresponding non-labeled sequence (Fig. 5f). This is comparable to the effect of a 50%–75% labeled tC^O^ transcript, which suggests that the external Cy5 modification indeed affects the ribosome translation capability more than tC^O^ does. We continued to monitor the mRNA-treated cells up to 72 h post-electroporation, and noted a continuous decrease in the GFP fluorescence signal over time with a 56%±5% loss for the unlabeled mRNA against 47%±8% loss for the tC^O^-labeled mRNAs when compared to the 24 h time point (Fig. 6a). This is likely a combined effect of mRNA degradation and cell division (the doubling time of SH-SY5Y cells under our experimental conditions is approximately 24 hours not appearing to be significantly impacted by tC^O^).

**Figure 6.**
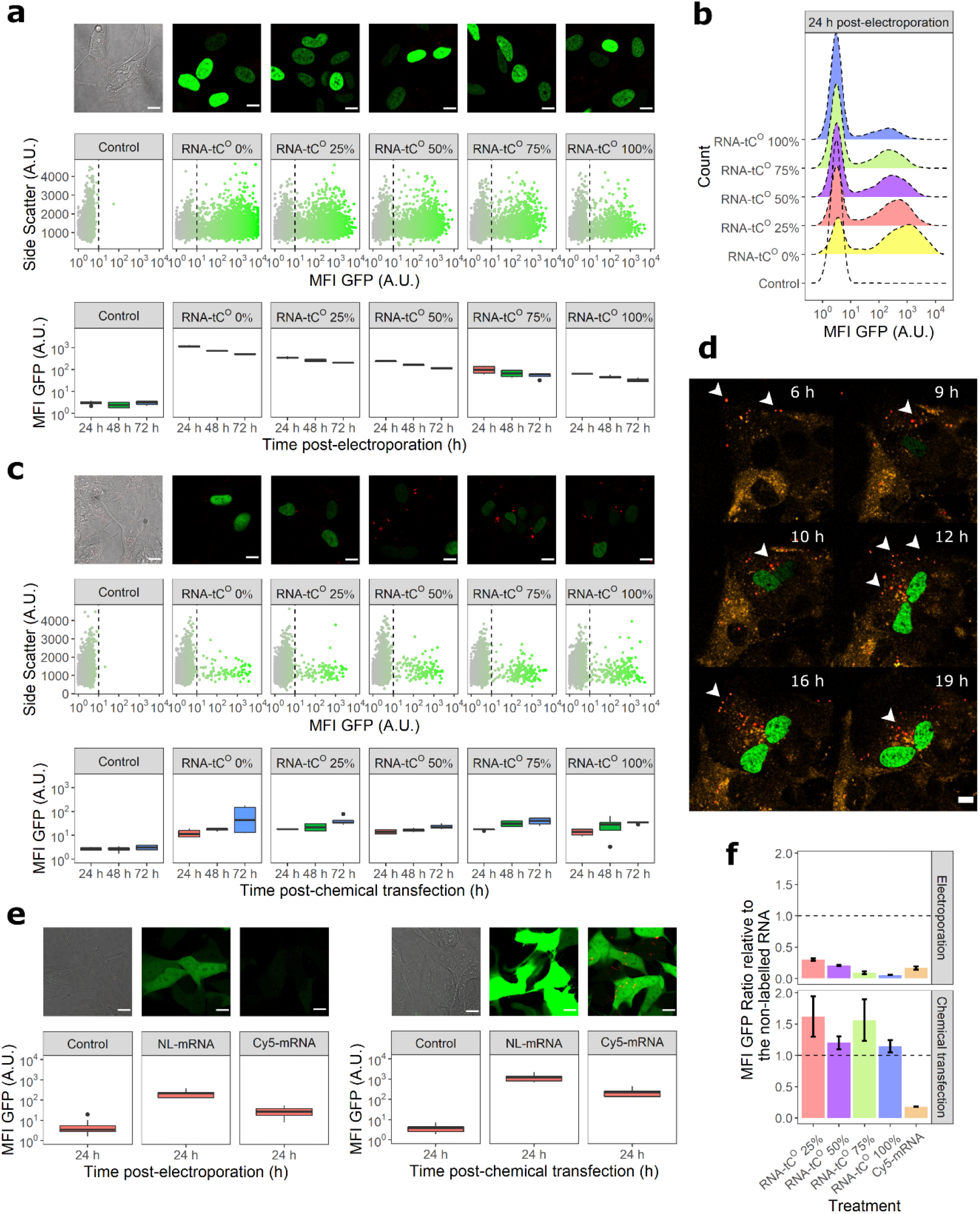
Translation efficiency of the modified RNA constructs in human cells and validation of tC^O^ as an intracellular tracking probe. The H2B:GFP encoded protein was observed by confocal microscopy and quantified by flow cytometry for each tC^O^-incorporated RNA constructs. Representative images (3x enlargements, scale bars: 10 μm), scatter plots, and histograms show the signal distribution in single living cells at (a, b) 24 h post-electroporation or (c) 48 h post-chemical transfection. The boxplots display the GFP mean (± standard deviation) fluorescence intensities (MFI GFP) up to 72 h from three independent experiments performed in triplicate. (d) Cells overexpressing mRFP-Rab5 (early endosome biomarker) were transfected with 75% tC^O^ mRNA and followed overtime to validate tC^O^ as an intracellular tracking probe (white arrows) not altering the translation, scale bars: 10 μm. (e) Cells were analyzed 24 h post-electroporation or post-transfection with non-labeled (NL) or Cyanine5-labeled (Cy5) eGFP encoding mRNAs (TriLink®), scale bars: 10 μm. (f) The impact of tC^O^ or Cy5 incorporation on RNA translation was expressed as the ratio of MFI GFP relative to the non-labeled RNA for all constructs.

It is evident from the images in Fig. 6a and 6e that neither the fluorescent tC^O^-labeled mRNA nor the Cy5-labeled mRNA can be detected inside cells after electroporation due to their low/diffused cytoplasmic concentration, as expected and in agreement with other reports.^65^ We therefore investigated the delivery of the H2B:GFP transcripts using a chemical transfection reagent (lipofectamine), which is also more relevant from a drug delivery perspective. This also resulted in the successful production of H2B:GFP (Fig. 6c and Supplementary Fig. 8b and 8d), demonstrating that the modified mRNA can undergo endosomal escape. The delivery efficiency was between 3.8%–4.6% of H2B:GFP-positive cells for the tC^O^-labeled transcripts compared to 12%-32% using electroporation. This reflects the relatively poor transfectability of SH-SY5Y cells compared to many other cell lines^66^ rather than the transfectability of tC^O^-labeled constructs, as further supported by the finding that cultures transfected with non-modified mRNA displayed a virtually identical response (4.5% of cells expressing H2B:GFP). An interesting point is that the H2B:GFP fluorescence increases gradually with time between 24 h and 72 h (Fig. 6c), which is a contrasting behavior compared to electroporated cells (Fig. 6a). This likely reflects the fact that the lipofectamine-mRNA complexes are continuously internalized and, potentially, that endocytosed complexes progressively release more transcripts with time, counteracting the degradation that mRNA experiences in the cytosol. The most noticeable finding of this experiment is that lipofectamine-delivered tC^O^-labeled mRNAs promote very similar H2B:GFP translation compared to the corresponding non-labeled mRNA, as indicated by the fluorescence levels in Fig. 6f. The Cy5-tagged mRNA, on the other hand, results in an average fluorescence level that is 80% lower than that of its corresponding non-labeled transcript. This suggests that tC^O^ does not impair the ability of mRNA to be processed by ribosomes upon chemical transfection, possibly because of an absence of charge and reduced steric hindrance, whereas Cy5, currently the most common commercial fluorophore for mRNA labeling, considerably impacts translation. It could also be an effect of the Cy5-labeling, as a result of its amphiphilic character, interacts differently with cellular membranes and/or delivery vehicle constituents and consequently impedes the transport of the Cy5-labeled mRNA to the location of translation.

Importantly, the complexation of the tC^O^-labeled mRNA with lipofectamine enabled its direct visualization inside cells using live cell confocal microscopy (Fig. 6c, red puncta). This represents the first observation of FBA-labeled nucleic acids inside live cells. We also showed that it is possible to visualize, in real time, both the uptake and subsequent translation of an FBA-modified mRNA by time-lapse recordings. We observed co-localization of the tC^O^ signal with an mRFP-labeled Rab5 protein, thus highlighting that the mRNA transits through the early endosome (Fig. 6d and Supplementary Movie 1). We were able to track both the intrinsically labeled mRNA transcripts and their translation products live, revealing spatiotemporal information on the delivery process, mRNA with as low as 50% tC^O^ content. This demonstrates the flexibility and versatility of this new labeling approach where fine-tuning of tC^O^ content can be utilized to optimize the mRNA for specific drug delivery applications and imaging needs.

## Conclusions

In this study we show how tC^O^, a fluorescent analogue of cytosine, in a remarkable way takes the role of natural cytosine and is correctly recognized by several enzymatic machineries, including the ribosome inside a human cell. We have developed a robust and affordable synthesis route of the tC^O^ triphosphate and, importantly, solid and straightforward methodologies to introduce it into RNA of different lengths, including full-length 1.2 kb mRNA, demonstrating that this base analogue is highly compatible with a variety of enzymatic processes. Furthermore, we demonstrate its suitability as an imaging-compatible label of RNA, for the first time realizing live cell imaging applications with a fluorescent base analogue. Overall, this clearly shows that the native properties of RNA are virtually unperturbed upon exchanging natural cytosine with our tC^O^.

Analysis of the in vitro transcription products, demonstrated that tC^O^ is incorporated into RNA virtually as efficiently as native CTP. Moreover, by sequencing PCR products stemming from the reverse transcription of modified RNA, we clearly demonstrated the high fidelity of an entire transcription-reverse transcription cycle starting with the incorporation of the modified nucleotide into RNA sequences and followed by conversion of the modified RNA to canonical DNA. We also demonstrated the modified transcripts to be non-toxic and translationally active both in bacterial lysate and in eukaryotic systems, regardless of their degree of tC^O^ incorporation. Finally, we showed how this conveniently allows for simultaneous monitoring of mRNA uptake and translation into H2B:GFP in live-cell confocal microscopy using selective excitation.

We envision that our straightforward approach for introducing non-perturbing fluorescent labels into RNA will be an excellent addition to existing imaging tools, applicable for elucidating trafficking mechanisms such as endosomal escape and exosome formation, both of which are of fundamental importance for pharmaceutical development. In principle, these methodologies can also be adapted to other base analogues (to access for instance different spectral properties), provided that the corresponding FBA triphosphate is tolerated by the enzymatic machinery as straightforwardly as the herein investigated tC^O^TP. Applying both our TdT end-labeling and T7 transcription strategies also holds significant combined potential as it would enable selective and site-specific incorporation of dual labels to allow for e.g. FRET applications. We are convinced that the development reported here will benefit pharmaceutical industry, clinical laboratories, and academic partners aiming at furthering their understanding of uptake and endosomal escape mechanisms and allow them to take vital steps towards new and improved delivery strategies for next-generation nucleic acid-based drugs.

## Materials and Methods

### Methods: Chemistry

Commercially available reagents were used without further purification. The following reagents used for the triphosphorylation were bought from Sigma-Aldrich: DCA deblock for ÄKTA, CAP A for ÄKTA, CAP B1 and B2 for ÄKTA, BTT Activator. ^1^H (500 MHz) and ^13^C (126 MHz) NMR spectra were recorded at 300 K on a Bruker 500 MHz system equipped with a CryoProbe. ^31^P (202 MHz) NMR spectra were recorded at 300 K on a Bruker 500 MHz system. All shifts are recorded in ppm relative to the deuterated solvent (DMSO-*d*_6_, CDCl_3_ or D_2_O).

#### 3-((2*R*,3*R*,4*S*,5*R*)-5-((bis(4-methoxyphenyl)(phenyl)methoxy)methyl)-3,4-dihydroxytetrahydrofuran-2-yl)-3*H*-benzo[*b*]pyrimido[4,5-*e*][1,4]oxazin-2(10*H*)-one 1

Compound was prepared according to the literature.^32^ MS (ESI^−^) [M-H]^−^=634.5. ^1^H NMR (500 MHz, DMSO-*d*_6_) δ 10.61 (bs, 1H), 7.42 (d, *J* = 7.7 Hz, 2H), 7.27 – 7.35 (m, 7H), 7.22 (t, *J* = 7.1 Hz, 1H), 6.90 (dd, *J* = 8.6, 4.2 Hz, 4H), 6.75 – 6.87 (m, 3H), 6.46 (d, *J* = 7.8 Hz, 1H), 5.71 (d, *J* = 3.6 Hz, 1H), 5.49 (bs, 1H), 5.18 (bs, 1H), 4.08 (d, *J* = 5.3 Hz, 1H), 4.04 (s, 1H), 3.94 (s, 1H), 3.71 (s, 3H), 3.70 (s, 3H), 3.29 (d, *J* = 4.8 Hz, 1H), 3.16 (d, *J* = 9.1 Hz, 1H).

Spectroscopic data in agreement with the literature.

#### Preparation of CPG solid support 3

The support **2** (1 g, 0.08 mmol) was activated by shaking in trichloroacetic acid 3% in DCE (8 mL, 0.08 mmol) for 18 h. The activated support was then filtered off and washed with 9:1 triethylamine:diisopropylethylamine (20 mL), dichloromethane (20 mL) and diethyl ether (20 mL). The activated support was dried under vacuum for 2 days before use.

Subsequently, the support (1 g, 0.08 mmol), succinic anhydride (0.345 g, 3.44 mmol) and *N*,*N*-dimethylpyridin-4-amine (0.070 g, 0.57 mmol) were suspended in dry Pyridine (3 mL) under N2. The reaction mixture was then gently shaken at RT for 4 h. After 4 h, solvent was filtered off and the support washed successively with pyridine (20 mL), dichloromethane (20 mL), diethyl ether (20 mL) and air-dried. Negative ninhydrin test on a small portion of support proved full succinylation.

Succinylated CPG could thereafter be kept at room temperature for several months.

#### CPG-supported (2*R*,3*R*,4*R*,5*R*)-5-((bis(4-methoxyphenyl)(phenyl)methoxy)methyl)-4-hydroxy-2-(2-oxo-2,10-dihydro-3*H*-benzo[*b*]pyrimido[4,5-*e*][1,4]oxazin-3-yl)tetrahydrofuran-3-yl acetate 4

In a 10 mL syringe with PTFE filter, succinylated support **3** (1.420 g, 82 μmol/g, 0.12 mmol), DMAP (0.028 g, 0.23 mmol), DIC (719 μl, 4.64 mmol), **1** (0.076 g, 0.12 mmol) and triethylamine (49 μl, 0.35 mmol) were suspended pyridine (5 mL). The mixture was gently shaken for 18 h at RT. After 18 h, the syringe was purged and the support washed with pyridine (5 mL), dichloromethane (5 mL) and diethyl ether. Subsequently, in the same syringe, DMAP (0.028 g, 0.23 mmol), DIC (719 μl, 4.64 mmol), triethylamine (49 μl, 0.35 mmol) and 2,3,4,5,6-pentachlorophenol (0.309 g, 1.16 mmol) were added to the support and suspended in pyridine (4 mL). The mixture was gently shaken for 4 h at RT before a solution of piperidine (2 mL, 20% in DMF – for capping of the unreacted carboxylic acids on the support) was added for 1 min (longer exposure time will reduce loading as piperidine cleaves the ester bonds with the nucleoside), then quickly washed away with DMF (3×5 mL), dichloromethane (5 mL) and diethyl ether (5 mL). Finally, the resin was shaken in a CAP A + CAP B mix (50/50 v/v) for 2 hours under argon atmosphere, then washed with DMF (5 mL), dichloromethane (5 mL), diethyl ether (5 mL) and argon-dried (final loading: 13 μmol/g – determined by reading optical density of a DMT solution cleaved from a weighed amount of support – ε=70000 M^−1^.cm^−1^ at 498 nm). Final loading can be increased by performing a second coupling with **1** in the same conditions before capping (typical loading after second coupling 15-20 μmol/g). Concentrating the reaction mixture and washing the residue multiple times with water and diethyl ether allows recovery of nearly 85 % of unreacted nucleoside **1**.

#### 6-chloro-*N*,*N*-diisopropyl-4*H*-benzo[*d*][1,3,2]dioxaphosphinin-2-amine 5

Compound was prepared according to the literature. ^67^ Briefly, 5-chlorosalicylic acid was reduced with LAH (0.5 equiv.) at −20 °C and the resulting 5-chlorosalicylic alcohol was cyclized into 2,6-dichloro-4*H*-benzo[*d*][1,3,2]dioxaphosphinine using PCl_3_ (1.2 equiv.) and triethylamine (2.3 equiv.) at −20 °C under argon. Low temperature and use of triethylamine as the base were decisive in avoiding rapid and quantitative Arbuzov rearrangement of the desired product into the more stable 2,5-dichloro-3*H*-benzo[*d*][1,2]oxaphosphole 2-oxide. The crude 2,6-dichloro-4*H*-benzo[*d*][1,3,2]dioxaphosphinine was subsquently treated with diisopropylamine (3 equiv.) for 2 h at room temperature. The mixture was then filtered under argon, concentrated to dryness and taken in 20 % diisopropylamine in heptane.

Quick filtration on a small silica gel plug allowed desired compound **5** as a colorless oil, crystallizing over time at −20 °C. Any attempt of more thorough column chromatography on compound **5** would lead to quantitative Arbuzov rearrangement. ^1^H NMR (500 MHz, DMSO-*d*_6_) δ = 7.23 (dd, *J* = 8.6, 2.6 Hz, 1H), 7.20 (d, *J* = 2.4 Hz, 1H), 6.92 (d, *J* = 8.6 Hz, 1H), 5.06 (dd, *J* = 14.7, 5.2 Hz, 1H), 4.89 (dd, *J* = 19.6, 14.8 Hz, 1H), 3.53 – 3.63 (m, 2H), 1.15 – 1.19 (dd, J = 8.0, 7.0 Hz, 12H).^31^P NMR (202 MHz, DMSO-*d*_6_) δ = 136.00 (s, 1P). Spectroscopic data in agreement with the literature.^49^

#### bis(tetrabutylammonium) dihydrogen diphosphate 6

Compound was prepared according to the literature.^49^ ^1^H NMR (500 MHz, D_2_O) δ 3.04 – 3.13 (m, 16H), 1.53 (bs, 16H), 1.24 (h, *J*=7.3, 16H), 0.83 (t, *J*=7.4, 24H). ^31^P NMR (202 MHz, D_2_O) δ = −10.78 (s, 2P). Spectroscopic data in agreement with the literature.

#### ((2*R*,3*S*,4*R*,5*R*)-3,4-dihydroxy-5-(2-oxo-2,10-dihydro-3H-benzo[*b*]pyrimido[4,5-*e*][1,4]oxazin-3-yl)tetrahydrofuran-2-yl)methyl triphosphate 7

Reaction was performed in a 5 mL syringe with PTFE filter loaded with **4** (800 mg, 0.016 mmol) under an argon atmosphere and shaking.

Steps were performed as following:

5’-DMT removal: the support was washed with a flow of DCA deblock until the filtrate was colorless, then washed with ACN (5 × 5 mL).

Coupling: *N*,*N*-diisopropyl-4*H*-benzo[*d*][1,3,2]dioxaphosphinin-2-amine **5** (345 mg, 1.36 mmol) was dissolved in 4.8 mL ACN and reacted portionwise with the support (3 equal couplings with reaction times 60 s – 60 s – 90 s respectively). To each coupling, BTT activator (2.4 mL) was also added. The support was subsequently washed with ACN (3×5 mL).

Oxidation: Pyridine/Water/Iodine (9/1/12.7 v/v/w, 5mL) for 45 s, followed by ACN wash (3×5 mL) and drying of the support in an argon flow.

Triphosphorylation: Two injections of bis(tetrabutylammonium) dihydrogen diphosphate 6 (0.5 M, 5 ml) for 15 min and 18 hours, respectively. The support was subsequently rinsed with DMF (5 mL), water (3×5 mL), ACN (5 mL) and then dried in an argon flow.

Cleavage and Purification: Cleavage of the triphosphate was done in 2 h at room temperature with AMA (50/50 v/v mix of 23 % aq. NH_4_OH and 40 % aq. methylamine, 5 mL). After 2 hours, the AMA filtrate was purged in a round-bottom flask and the support was rinsed 3 times with 23% aq. NH_4_OH solution. After freeze-drying of the mixture, purification by HPLC (Waters Acquity HSS T3 column, 2.1×50 mm, 0.4 mL/min, 2 to 99 % 50 mM NH_4_OAc in water 80:20 EtOH) was performed to allow compound **7** (5.6 mg, 62.0 % determined from UV absorbance) as a light-yellow solid (ammonium salt). The same level of purity could be achieved with ion-exchange HPLC using a semi-preparative Dionex DNAPac PA100 column (9 × 250 mm) on an ÄKTA pure 25 HPLC system using a gradient from water to 20 % 1M NH_4_HCO_4_ (pH 7.8) in 30 min at a flow rate of 4 mL/min.

HRMS (ESI-TOF) *m/z* calc. for C_15_H_18_N_3_O_15_P_3_ [M+H]^+^: 574.0029, found : 574.0013; *m/z* calc. for C_15_H_18_N_3_O_15_P_3_ [M-H]^−^:571.9878, found: 571.9872.

^1^H NMR (500 MHz, D_2_O) δ 7.44 (s, 1H), 6.84 – 6.94 (m, 3H), 6.79 (dd, *J* = 7.5, 1.7 Hz, 1H), 5.91 (d, *J* = 4.9 Hz, 1H), 4.36 (t, *J* = 4.8 Hz, 1H), 4.29 (t, *J* = 5.1 Hz, 1H), 4.25 (d, *J* = 4.1 Hz, 3H). ^13^C NMR (126 MHz, D_2_O) δ 155.8, 154.8, 142.4, 129.4, 124.9, 124.3, 122.3, 116.6, 88.8, 82.8, 73.4, 69.7, 64.5. ^31^P NMR (202 MHz, D_2_O) δ −10.89 (d, *J* = 18.5 Hz, 1P), −11.46 (d, *J* = 19.7 Hz, 1P), −23.21 (t, *J* = 19 Hz, 1P).

### Methods: Spectroscopy

The tC^O^-RNA products from the cell-free transcription reactions (prior to polyadenylation and capping, see Methods: Bio for details) were measured as received, *i.e.* in RNAse free Milli Q water. All measurements were carried out at room temperature (*ca.* 22°C) in a 3.0 mm path length quartz cuvette, with a sample volume of *ca.* 60 μL.

#### Steady state absorption

Absorption spectra were recorded on a Cary 5000 (Varian Technologies) spectrophotometer with a wavelength interval of 1.0 nm, integration time of 0.1 s, and a spectral band width (SBW) of 1 nm. All spectra were baseline corrected by subtracting the corresponding absorption from the solvent only. A second-order polynomial Savitzky-Golay (five points) smoothing filter was applied to all spectra. For samples exhibiting significant scattering, as evidenced by characteristic absorption in the long wavelength region (here for λ > 475 nm), an additional correction was applied. The scattering contribution (*A*_*scatter*_) to the absorption was in such cases fitted (using absorption at 550–475 nm as input) to the Rayleigh scattering function (equation S1), where *c* is a proportionality constant and *A*_0_ a constant, and then subtracted for all wavelengths.

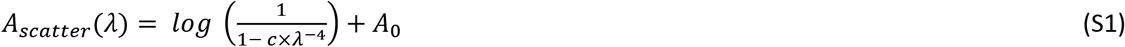

#### Steady state emission

Emission spectra were recorded on a SPEX Fluorolog (Jobin Yvon Horiba) fluorimeter with excitation at 356 nm. Emission was collected at a right angle with an integration time of 0.1 s and wavelength interval of 1 nm. Monochromator slits were adjusted to achieve optimal signal output, leading to SBWs in the interval 1.5–2.5 nm on both the excitation and emission side.

Emission spectra were corrected for Raman scattering from water by subtracting the corresponding emission from a sample containing only solvent. A second-order polynomial Savitzky-Golay (five points) smoothing filter was applied to all spectra.

#### Fluorescence quantum yield determination

Sample fluorescence quantum yields (*Φ*_*F*_) were determined relative to a solution of quinine sulphate (Sigma) in 0.5 M H_2_SO_4_ (*Φ*_*F*,*REF*_ = 0.546)^68^ and calculated according to equation S2.

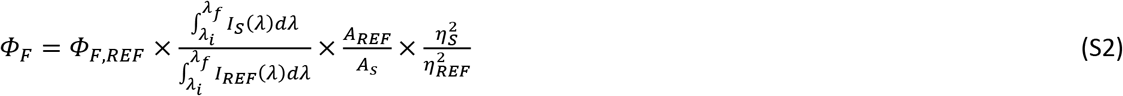

Emission spectra for the sample, *I*_*S*_(*λ*), and reference, *I*_*REF*_ (*λ*), were integrated between *λ*_*i*_ = 365 nm and *λ*_*f*_ = 700 nm. Absorption at the excitation wavelength (356 nm) for the sample (*A*_*s*_) and reference (*A*_*REF*_) were in the interval 0.05–0.11 for all samples. Adopted solvent refractive indices for the samples (water) and reference (0.5 M H_2_SO_4_) were *η*_*S*_ = 1.333 and *η*_*REF*_ = 1.339, respectively. All quantum yields are presented as mean ± standard deviation of two independent cell-free transcription reactions.

#### Time-resolved emission

Fluorescence lifetimes were determined using time-correlated single photon counting (TCSPC). Samples were excited using an LDH-P-C-375 (PicoQuant) pulsed laser diode with emission centered at 377 nm (FWHM pulse width was 1 nm and 70 ps with respect to wavelength and time, respectively), operated with a PDL 800-B (PicoQuant) laser driver at a repetition frequency of 10 MHz. Sample emission (458 nm, SBW = 10 nm) was collected at a right angle, through an emission polarizer set at 54.9° (magic angle detection). Photon counts were recorded on a R3809U-50 microchannel plate PMT (Hamamatsu) and fed into a LifeSpec multichannel analyzer (Edinburgh Analytical Instruments) with 2048 active channels (24.4 ps/channel), until the stop condition of 10^4^ counts in the top channel was met. The instrument response function (IRF) was determined using a frosted glass (scattering) modular insert while observing the emission at 377 nm (SBW = 10 nm).

#### Fitting of fluorescence lifetimes

The intensity decays were fitted with IRF re-convolution to the multiexponential model shown in equation S3.

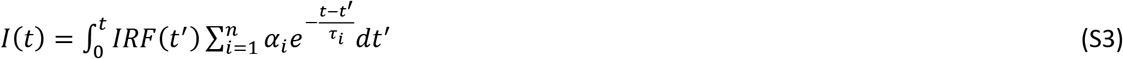

The least-square re-convolution fitting procedure was carried out using the DecayFit software (http://www.fluortools.com/software/decayfit). All decays were fitted to a tri-exponential (*n* = 3) model. The presented lifetimes are amplitude-weighted average lifetimes 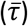, calculated using the pre-exponential factors (*α*_*i*_) and lifetimes (*τ*_*i*_) according to equation S4. The fitting parameters for the decays are shown in Supplementary Table 2.

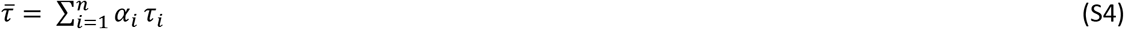

#### Cell-free transcription reaction kinetics

The ratio of the rate constants for cytosine *vs.* tC^O^ incorporation 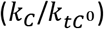 was calculated using the absorption spectra of the tC^O^-RNA transcripts (*A*_260_ and *A*_369_), and triphosphate initial concentrations ([*CTP*]_0_ and [*tC*^0^*TP*]_0_) as input. Equations S5 and S6 follows upon assuming first-order reaction kinetics with respect to the triphosphate species [CTP] and [tC^O^TP].

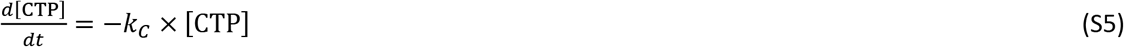

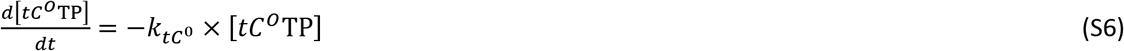

Solving S5 and S6 for the respective rate constants renders equation S7, in which [C] and [tC^O^] denote the concentration of *incorporated* C and tC^O^, respectively.

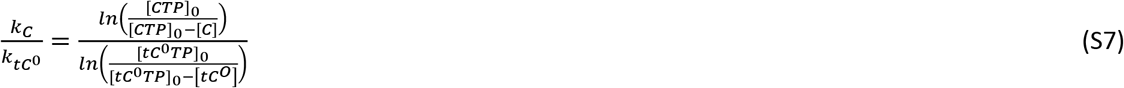

Using the Beer-Lambert law, absorption is related to nucleobase concentration according to equations S8 and S9. The following molar absorptivities (unit: M^−1^cm^−1^) were adopted: 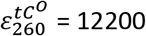, 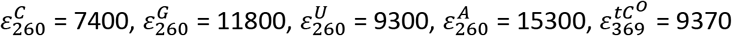. It should be noted that the molar absorptivity of the tC^O^ adopted here 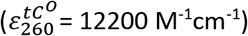 differs slightly from what has been published previously by our group (11000 M^−1^cm^−1^). This is due to a gravimetric re-evaluation of this parameter using a large amount of compound prior to starting this work.

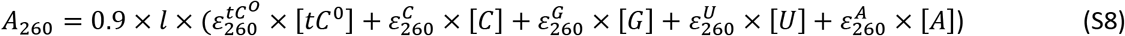

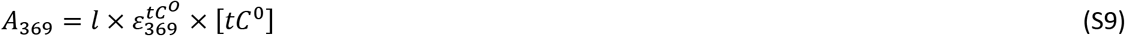

Assuming that the product RNA is uniform in size (1247 nucleotides), its base composition (A: 408, U: 272, G: 307, C: 260) allows for equations S10 – S13.

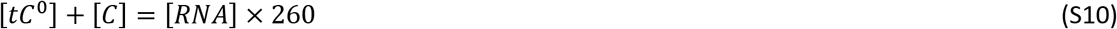

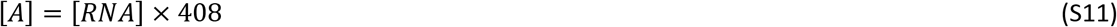

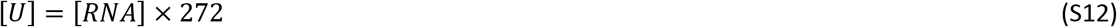

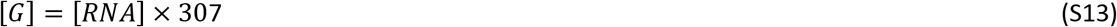

Solving the equation system composed of S7 through S13 allows for quantification of 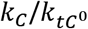, [*tC*^0^], [*RNA*], [*C*], [*A*], [*U*], and [*G*]. The average-strand *tC*^*0*^ incorporation degree 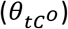 can then be calculated according to equation S14.

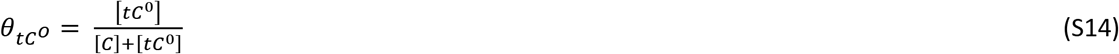

Using the volume of the cell-free reaction (50 μL) and resulting product solution (100 μL), equation S15 was applied to calculate the *tC*^0^ incorporation yield 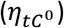.

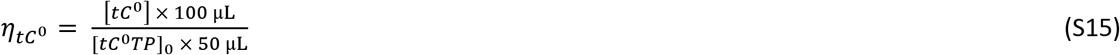

Consonantly, the RNA yield (*η*_*RNA*_) was calculated according to equation S16.

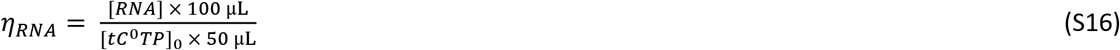

### Methods: Biochemistry

The TdT, Taq, and Phusion polymerases were purchased from New England Biolabs as well as the natural dNTPs and NTPs. The AmpliScribe™ T7 High Yield Transcription Kit was purchased from Epicentre and the HiScribe T7 In Vitro Transcription Kit was obtained from New England Biolabs. The reverse transcriptase M-MLV RT as well as the T4 DNA ligase were obtained from Promega. The pGEM-T Vector kit was purchased from Promega. Acrylamide/bisacrylamide (29:1, 40 %) was obtained from Fisher Scientific. Visualization of PAGE gels was performed by fluorescence imaging using a Storm 860 or a Typhoon Trio phosphorimager with the ImageQuant software (GE Healthcare). Natural DNA and RNA oligonucleotides were purchased from Microsynth. Concentrations of short oligonucleotides were quantitated by UV spectroscopy using a UV5Nano spectrophotometer (Mettler Toledo).

#### General protocol for the TdT-mediated polymerization reactions

A solution containing 20 or 2 pmol of the fluorescently labeled ssRNA oligonucleotides (TdT1, TdT2, and TdT3), 1 U/μL of TdT, the modified nucleotide (200 μM final concentration), 1x reaction buffer (from Promega (containing 1 mM Co^2+^) for Supplementary Fig. 1a and NEB for Supplementary Fig. 1b), metal cofactor (0.25 mM Co^2+^, 1 mM Mn^2+^, or 1 mM Mg^2+^ (final concentrations)) and H_2_O (for a total reaction volume of 10 μL) was incubated at 37 °C for different reaction times. The reactions were then quenched by addition of 10 μL of loading buffer. The reaction products were then resolved by electrophoresis (PAGE 20 %) and visualized by phosphorimager analysis. Alternatively, products stemming from the TdT-labeling of oligonucleotide TdT3 were irradiated with UV light prior and after purification with a NucleoSpin (Macherey-Nagel) clean-up kit.

#### In vitro transcription from dsDNA templates

**Buffer compositions**:

1. buffer supplied from Epicentre in the AmpliScribe T7 High Yield Transcription Kit.
2. 40 mM Tris-acetate pH 8.0, 10 mM Mg(OAc)_2_, 0.5 mM MnCl_2_, 5 mM DTT, 0.1% Tween 20, 8 mM spermidine.^69^
3. 40 mM Tris·HCl, pH 8.0, 30 mM MgCl_2_, 3.75 mM MnCl_2_, 20 mM DTT, 0.01% Triton X-100, 3.6 mM spermidine.^70^
4. 40 mM Tris·HCl pH 8, 12 mM MgCl_2_, 5 mM DTT, 0.01% Triton-X, 1 mM spermidine, 4% glycerol.^56^
5. 200 mM HEPES pH 7.5, 5.5 mM MgCl_2_, 2 mM spermidine, 40 mM DTT, 0.01% Triton, 10% PEG8000, 1.5 mM MnCl_2_, 10 U/ml yeast inorganic pyrophosphatase.^71^
6. 40 mM Tris·HCl pH 8, 8 mM MgCl_2_, 2 mM spermidine, 25 mM NaCl.^35^

#### i) Transcription reaction with short dsDNA templates

The T7 promoter (oligonucleotide T2) was hybridized with one of the DNA templates (B2, DNA3, DNA4) by heating at 75 °C for 3 min and then cooling down for 10 min on ice. The hybridized oligonucleotides were then added to a mixture containing 0.4 mM of each rNTP, RNAse inhibitor, DTT (10 mM final concentration for reactions with the AmpliScribe kit), reaction buffer, and RNAse free water and lastly 1.5 μL of T7 RNA polymerase (HiScribe T7 In Vitro Transcription Kit from New England Biolabs or AmpliScribe T7 High Yield Transcription Kit from Epicenter). The reaction mixtures were incubated for 1 h (Supplementary Fig. 2a) or 0.5 h (Supplementary Fig. 2b) at 37 °C and the reactions were quenched by adding 2 μL EDTA 100 mM. The transcription products were precipitated by EtOH and then analyzed by gel electrophoresis (PAGE 20%) and imaged after staining with Midori green or silver.

#### ii) Transcription reactions with longer dsDNA templates

DNA duplexes were prepared by PCR either using templates T12.3 or D1 Library^72^ along with primers FWD-D1-RNA 2 and REV-D1-RNA 2. The PCR reaction mixture with template T12.3 consisted of 0.2 mM dNTPs, 1 μM of each primer, 2.6 ng/μL of template, 1x Ex Taq reaction buffer, and 20 U of Ex Taq DNA polymerase in a total volume of 800 μL. The PCR program started with a 5 min long denaturation step (95 °C) and was followed by 29 PCR cycles: 1. 30 sec at 95 °C, 2. 30 sec at 62 °C, 3. 1 min at 72 °C and a final elongation step (10 min at 72 °C). The PCR reaction mixture for the preparation of the dsDNA library consisted of 0.2 mM dNTPs, 1 μM of each primer, 72 nM template, 1x Thermopol reaction buffer, 1.5 mM MgCl_2_, 0.5 mg/mL BSA, and 20 U of Taq DNA polymerase in a total volume of 800 μL. The PCR program started with a 5 min long denaturation step (95 °C) and was followed by 5 PCR cycles: 1. 1 min at 95 °C, 2. 1 min at 54 °C, 3. 1.5 min at 72 °C and a final elongation step (10 min at 72 °C). All PCR products were then purified by phenol/chloroform extraction followed by ethanol precipitation and filtration (10K Nanosep). All transcription reactions proceeded as described in section i) using reaction buffer 5. Gel analyses of the transcription reactions are highlighted in the manuscript (for D1 Library) and Supplementary Fig. 3 (for T12.3; see following paragraph).

### Cloning and sequencing (RT-PCR stemming from template T12.3)

The products stemming from transcription reactions (both 30 min and 6 h reaction times) with CTP or **tC°TP** were reverse transcribed and then cloned and sequenced to assess the incorporation fidelity of the modified nucleotides during both transcription and reverse transcription.

#### i) reverse transcription

20 pmol of the products (modified or natural) stemming from in vitro transcription reactions (see Section 4 and transcription reactions using template T12.3) are added to a reaction mixture containing 200 pmol primer REV-D1 128, the mixture is heated to 70 °C for 5 minutes then kept on ice for 10 seconds, then 0.2 mM dNTPs, RT reaction buffer, and 200 U of reverse transcriptase M-MLV RT are added in 50 μL final volume. The resulting reaction mixtures were then incubated at 42 °C for 1 hour. The resulting reverse transcripts were then purified by 10K Nanosep columns and DNA was quantified by UV absorbance. Gel analysis of products stemming from the transcription and reverse transcription reactions is shown in Supplementary Fig. 3.

#### ii) PCR amplification and A-tailing

PCR reaction mixtures containing 0.1 pmol of the reverse transcription products, 0.5 μM primers **REV_D1 23** and **REV-D1 128**, 0.2 mM dNTPs, 0.02 U/μL of Phusion DNA polymerase, and Phusion reaction buffer were prepared for a total volume of 300 μL. After an initial denaturation step (98 °C for 30 seconds), 25 PCR cycles were carried out using the following program: 1. 98 °C for 10 seconds; 2. 54 °C for 30 seconds; 3. 72 °C for 20 seconds. A 5-minute final elongation step was also included in the program. After PCR, the reaction mixtures were purified using MiniElute purification kits (Qiagen) and subjected to A-tailing reactions. Briefly, 5 μL of the purified PCR mixtures were incubated for 30 min at 70°C in the presence of 1.5 mM MgCl_2_, 0.2 mM ATP, 5U of Taq DNA polymerase, and reaction buffer 1x.

#### iii) Ligations and transformations

Ligation reactions were carried out by preparing a reaction mixture containing 1 μL of the A-tailed PCR reaction products, 1 μL of the pGEM T vector (50 ng), T4 DNA ligase (6 Weiss U), and ligase reaction buffer in a total volume of 11 μL. The reaction mixtures were then incubated at room temperature under gentle stirring for 1 h. The resulting ligation products were then purified by filter dialysis (nitrocellulose 0.05 μM, Millipore). 3 μL of the ligation reactions were then added to 50 μL of beta 2033 competent cells (frozen instead of freshly prepared). The cells were then subjected to electroporation at 1800V. After addition of 1 mL LB broth, the cells were incubated at 37 °C for 15 min. 100 μL of the mixtures were then streaked onto Petri dishes and centrifuged for 3 min at 8000 rpm. The supernatants were then removed and the resuspended pellets were streak onto LB-Carbenicillin (100 μg/ml final) / X-Gal (80 μg/mL) / IPTG (0.5 mM final) plates. The plates were then incubated at 37 °C for 12 h and white colonies were selected and grown overnight at 37 °C in LB containing carbenicillin medium (100 μg/mL), and plasmids were purified by use of a QIAprep Spin Miniprep Kit according to the manufacturer’s instructions (QIAGEN). The selected plasmids were purified by use of a QIAprep Spin Miniprep Kit according to the manufacturer’s instructions (QIAGEN). For the reaction products stemming from transcription with natural CTP, 41 plasmids were purified for the 30 min reaction products and 39 for the 6 h reaction time point. For the products with the modified triphosphate, 38 plasmids were purified for the 30 min reaction products and 39 for the 6 h reaction time point.

#### iv) Sequencing and analysis

The sequencing (Illumina) of the purified plasmids was carried out using the Eurofins sequencing service with the sequencing primers M13 uni (−43) and M13 rev (−29) for the PCR amplification of the plasmids. Sequence alignment was performed with Benchling [Biology Software] (2018) using MAFFT Algorithm (https://benchling.com). Motif logos highlighted in the manuscript were made from these alignments using Weblogo 2.0.^73^ The sequence alignments obtained (only shown for the products obtained with primer M13 uni (−43)) are shown in Supplementary Fig. 4 and 5.

### Generation of H2B:GFP DNA template

The original coding sequence for H2B:GFP was taken from pCS2-H2B:GFP plasmid (Addgene, Plasmid #53744, Supplementary Fig. 6), manually codon-optimized to minimize the occurrence of poly-C_n_ stretches (n<3), in silico-assembled with an additional T7 promoter and other desired features (Shine-Dalgarno/Kozak consensus sequences for enhancement of translation and a 3xStop, respectively at the 5′ and 3′ of the coding sequence itself, plus the needed HindIII/SnaBI restriction sites, to generate the ligation-prone sticky ends) and ordered from Twist Bioscience as a synthetic gene block (Supplementary Fig. 6). The obtained sequence was then PCR-amplified, using a Phusion Hot Start High-Fidelity Taq (Thermo Scientific), and subcloned into a HindIII/SnaBI-digested (Fast Digest enzymes, Thermo Scientific) empty pCS2 backbone. After ligation with T4 ligase for 1 h at room temperature (Roche), DH5α E. coli competent cells (Invitrogen) were transformed following the recommended protocol, and obtained colonies were screened by colony-PCR. The selected colony was then inoculated into a midiprep-scale volume of liquid Luria-Bertani growth medium (VWR) and plasmid DNA isolated using a PureLink Fast Low-Endotoxin Midi Plasmid Purification Kit (Thermo Scientific). The purified plasmid was finally digested again with HindIII/SnaBI and gel-purified, to generate the transcription template with the desired size.

Primers (Eurofins Genomics):

Twist-H2B.F: GAAGTGCCATTCCGCCTGAC
Twist-H2B.R: CACTGAGCCTCCACCTAGCC

### H2B:GFP RNA transcription and purification

In vitro transcription reactions, for T7 and SP6 polymerases (Thermo Scientific), were assembled as recommended by the corresponding protocols, with few modifications that resulted in a consistently increased yield in all conditions:

5X Transcription buffer − 10 μl
NTP Mix, 10 mM each (2 mM final concentration) – volume depending on batch for tC^O^TP
Linearized template DNA 1 μg – volume depending on concentration
RiboLock RNase Inhibitor − 1.25 μl (50 U)
T7/T3/SP6 RNA Polymerase − 3 μl (60 U, double compared to recommendations)
MgCl_2_ 4 mM final concentration (increased as recommended by Thomen et al.)^74^
DEPC-treated Water qsp 50 μl

In vitro transcriptions were always performed at 20 °C for 14 h, then RNAs were purified using a Monarch RNA Cleanup kit (NEB), or homemade equivalent buffers and regenerated columns,^75^ following the same rationale. It was possible to partially recover unreacted tC^O^TP from the transcription mixtures by HPLC to re-use for further assays. For cellular studies, each batch of RNA was then enzymatically added with a polyA tail (with a Poly(A) Polymerase, NEB protocol #M0276 with incubation extended to 1 hour) and a Cap 0 analog (using a Vaccinia capping system, NEB protocol #M2080), following the recommended procedures. This was necessary to increase, respectively, affinity for the cellular translation machinery and resistance to endogenous nucleases.

### Denaturing bleach-agarose gels

For a qualitative check of all in vitro synthesized RNAs, a denaturing agarose gel was run, in presence of 1,5 % bleach (Sigma Aldrich), as recommended in Aranda et al.^76^ RNAs were first mixed with a 6x DNA loading dye (Invitrogen) and then heat-denatured at 70°C for 10 min in a heating block, then immediately transferred and kept on ice. The RiboRuler High Range RNA Ladder (Thermo Scientific) underwent the same treatment; 2 μl of RNA ladder were loaded along the samples and the gel was run at constant voltage (70 V) for 1 h and then imaged, under UV transillumination (302 nm) using a ChemiDoc Touch (BioRad). To counterstain the whole gel, and especially the lanes without tC^O^TP-containing samples, a standard ethidium bromide staining was finally performed at room temperature for 10 min and gentle rocking, followed by two washes in TAE and then a final wash in distilled water (10 min each).

### Cell culture

Human neuroblastoma SH-SY5Y cells (Sigma-Aldrich) were grown in a 1:1 mixture of minimal essential medium (HyClone) and nutrient mixture F-12 Ham (Sigma-Aldrich) supplemented with 10% fetal bovine serum (FBS), 1% non-essential amino acids (Lonza) and 2 mM L-glutamine. For the tracking experiments, an in-house generated model of human hepatic Huh-7 cells stably overexpressing mRFP-Rab5 were cultured in DMEM/GlutaMax/High glucose (Gibco) supplemented with 10 % FBS. The cells are detached with trypsin-EDTA 0.05 % (Gibco) and passaged twice a week.

### Electroporation or chemical transfection

Cells were electroporated either with 9.7 μg of tC^O^TP (for in vitro incorporation experiments) or 100 ng of tC^O^-labeled mRNA per 10^5^ cells (for in vitro translation, cytotoxicity assessment, flow cytometry analysis and confocal microscopy), using a Neon Transfection System (Invitrogen, Carlsbad, CA, US) and following the protocol for 10 μL Neon Tip provided by the manufacturer, with a triple pulse of 1200 V and a pulse width of 20 ms. For chemical transfection, SH-SY5Y cells were seeded one day prior transfection at a density of 0.8 10^5^ cells/mL, in 48-well plate or glass-bottomed culture dishes for flow cytometry of confocal microscopy analysis, respectively. Lipofectamine MessengerMAX was used as chemical reagent for transfection according to the manufacturer’s instructions. Briefly, the reagent was diluted and incubated for 10 min at room temperature in Opti-MEM medium. The tC^O^-mRNA constructs were added to the reagent to reach a 1:1 final ratio reagent-mRNA (v/w), followed by a 5 min incubation at room temperature allowing the complex mRNA-lipid to form. Cells were incubated with this complex up to 72 h. To address the impact of the dye incorporation on RNA translation, SH-SY5Y cells were electroporated or chemical transfected with commercially available non-labeled (NL) or Cyanine5-labeled (Cy5) eGFP encoding mRNAs (Trilink®) has described here.

### Cytotoxicity assessment

Cell membrane integrity was determined using the Pierce™ LDH Cytotoxicity Assay Kit (Invitrogen) according to the manufacturer’s instructions. Briefly, LDH released in the supernatants of cells 24 h post-electroporated or post-transfected with tC^O^-labeled mRNA, or Cy5-mRNA, was measured with a coupled enzymatic assay which results in the conversion of a tetrazolium salt into a red formazan product. The absorbance was recorded at 490 nm and 680 nm. The toxicity was expressed as the percentage of LDH release in supernatant compared to maximum LDH release (supernatant + cell lysate). Data are means ± SD from three experiments performed in triplicate.

### Flow cytometry

Following electroporation of tC^O^-labeled mRNA, cells were seeded in 48-well plate (2.10^5^ cells/well) and the expression of H2B:GFP in cells was quantified by flow cytometry. Briefly, 24 h, 48 h or 72 h post-electroporation or post-transfection with tC^O^-mRNA, non-labeled mRNA or Cy5-mRNA, cells were harvested and analyzed on a Guava EasyCyte 8HT flow cytometer (Millipore). Data are mean fluorescence intensities ± SD of gated single living cells from three experiments performed in triplicate. The average fluorescence intensities were baseline corrected by subtracting the signal for RNase-free water electroporated or transfected cells. All flow cytometry data were analyzed in Flowing software (version 2.5.1) and displayed using R (http://www.R-project.org/).^77^

H2B:GFP: Excitation 488 nm; Emission 525-530 nm.

### Confocal microscopy

After electroporation, cells were seeded in glass-bottomed culture dishes (MatTek glass-bottomed or in 4-sectors subdivided CELLview dishes; 2.10^5^ cells/chamber). For tracking experiment, the Huh-7 cells stably overexpressing mRFP-Rab5 were incubated with lipofectamine/tC^O^-mRNA complex and time-lapse was recorded up to 20 h post-chemical transfection. Confocal images were acquired on a Nikon C2+ confocal microscope equipped with a C2-DUVB GaAsP Detector Unit and using an oil-immersion 60 × 1.4 Nikon APO objective (Nikon Instruments, Amsterdam, Netherlands). Data were processed with the Fiji software.^78^

H2B:GFP: Excitation 488 nm; Emission 495-558 nm.
tCO-labeled mRNA: Exc. 405 nm; Em. 447-486 nm.
Cy5-labeled mRNA: Exc. 640 nm; Em. 652-700 nm.
mRFP-Rab5: Exc. 561 nm; Em. 565-720 nm.

### Cell-free translation

Cell-free translation reactions were performed using E. coli bacterial lysates and an Expressway™ Mini Cell-Free Expression System (Thermo Scientific). Calmodulin-like 3 protein is provided as a positive control plasmid (pEXP5-NT/CALML3) in the kit itself; this DNA vector contains a T7 polymerase promoter and a 6xHis tag, hence it was first in vitro-transcribed in presence of the desired concentrations of tC^O^TP (*vide supra*). The obtained RNAs, once purified, were used as templates for the cell-free translation reaction according to the manufacturer’s recommendations:

E. coli slyD-Extract − 20 μl
2.5X IVPS E. coli Reaction Buffer (−A.A.) − 20 μl
50 mM Amino Acids (−Met) − 1.25 μl
75 mM Methionine* − 1 μl
T7 Enzyme Mix − 1 μl (omitted when using tC^O^-labeled RNAs)
DNA Template − 1 μg (when testing the tC^O^-labeled RNAs, added the same amount of RNA instead)
DNase/RNase-free distilled water qsp 50 μl

### Coomassie staining and Western Blots

Protein samples from in vitro translation experiments were quantified with a Qubit Protein Assay kit (Thermo Scientific), mixed with 6x SDS Laemmli reducing buffer (Alfa Aesar), then heat-denatured at 85 °C for 10 min and kept at room temperature until needed. Samples were generally run in 1 mm polyacrylamide 4-20 % Novex MES/SDS gels (Thermo Scientific) and using a Mini Gel Tank, with the PSU set at constant voltage (200 V). For Coomassie staining, the gel was then washed three times in boiling water, to remove excess of SDS, on a benchtop shaker; a 1x Coomassie non-toxic staining solution was added to the gel and microwaved until initial boiling.^79^ Gel was finally washed after the appropriate incubation time, to remove excess of background noise, in distilled water and imaged using a ChemiDoc Touch. For Western Blot, the gels were blotted onto PVDF LF ethanol-activated membranes (BioRad) with a TransBlot semi-dry system (BioRad), according to manufacturer’s recommendations (settings for 1mm-thick gels and mixed weight proteins). PVDF membranes were then washed 5 min in TBS-T (TBS and 0,1 % Tween-20, Sigma Aldrich), blocked in 5 % milk in TBS for 1 h at room temperature and incubated with the appropriate primary antibody dilutions. After 3×5 min washes in TBST and an incubation of 1 h with the corresponding HRP-conjugated secondary antibodies, the membrane was washed again three times in TBS-T, once in TBS and once more in distilled water. Finally, membranes were incubated with a minimal volume of SuperSignal West Pico PLUS (Thermo Scientific) and imaged with a ChemiDoc Touch. Primary antibodies: mouse monoclonal anti-6xHistidine tag (Invitrogen) and mouse monoclonal anti-GAPDH (ref. 437000, Invitrogen), both diluted 1:1000 in 3 % BSA/TBS-T. Secondary antibodies: HRP-conjugated polyclonal goat anti-Ms and anti-Rb Cross-Adsorbed IgG (H+L) (ref. A16072 and A16104, Invitrogen), used at 1:10000 dilution in TBS-T.

## Data availability

The data that support the plots within this paper and the findings of this study are available from the corresponding author upon reasonable request.

## Acknowledgements

We gratefully acknowledge Prof. Tom Brown at the University of Oxford for fruitful discussions about the triphosphate synthesis and Linda Thunberg at AstraZeneca Gothenburg for help with purification of the triphosphate. This work was conducted within the FoRmulaEx research consortium and with associated financial support to E.K.K and L.M.W from the Swedish Foundation for Strategic Research (SSF, grant No. IRC15-0065). F. L.-A., I. S. and M. H. acknowledge funding from Institut Pasteur.

## Contributions

L.M.W. and E.K.E. conceived the idea. L.M.W, E.K.E, M.H. and A.D. designed and supervised the study. T.B. and I.S. designed and performed the synthesis of the modified triphosphate. E.C. and C.G. did the in vitro transcription assays and E.C. the in vitro translation assays. C.G. performed the reverse transcriptions and sequencing. F.L.-A. did the TdT labeling assays. J.R.N. designed and implemented the photophysical study and determination of the incorporation rates during transcription. A.G. setup and performed the flow cytometry, imaging and cytotoxicity analysis for the cell experiments. T.B. was main responsible for writing the manuscript whereas E.K.E. and L.M.W. made major contributions to the writing process. All authors contributed to revising the draft.

## References

1 Crooke, S. T., Witztum, J. L., Bennett, C. F. & Baker, B. F. RNA-Targeted Therapeutics. Cell Metabolism 27, 714–739 (2018).

2 Dowdy, S. F. Overcoming cellular barriers for RNA therapeutics. Nat. Biotechnol. 35, 222–229 (2017).

3 Conner, S. D. & Schmid, S. L. Regulated portals of entry into the cell. Nature 422, 37 (2003).

4 Crooke, S. T. et al. Cellular uptake and trafficking of antisense oligonucleotides. Nat. Biotechnol. 35, 230 (2017).

5 Pei, D. & Buyanova, M. Overcoming Endosomal Entrapment in Drug Delivery. Bioconjugate Chem. (2018).

6 Shimomura, O., Johnson, F. H. & Saiga, Y. Extraction, Purification and Properties of Aequorin, a Bioluminescent Protein from the Luminous Hydromedusan, Aequorea. J. Cell. Compar. Physl. 59, 223–239 (1962).

7 Jung, G. in Fluorescent Analogues of Biomolecular Building Blocks: Design and Applications Ch. 4, 55–90 (Wiley, 2016).

8 Wittrup, A. et al. Visualizing lipid-formulated siRNA release from endosomes and target gene knockdown. Nat. Biotechnol. 33, 870–876 (2015).

9 Gaus, H. J. et al. Characterization of the interactions of chemically-modified therapeutic nucleic acids with plasma proteins using a fluorescence polarization assay. Nucleic Acids Res. 47, 1110–1122 (2019).

10 Hughes, L. D., Rawle, R. J. & Boxer, S. G. Choose Your Label Wisely: Water-Soluble Fluorophores Often Interact with Lipid Bilayers. PLOS ONE 9, e87649 (2014).

11 Li, Y., Ke, K. & Spitale, R. C. Biochemical Methods To Image and Analyze RNA Localization: From One to Many. Biochemistry 58, 379–386 (2019).

12 Burke, K. S., Antilla, K. A. & Tirrell, D. A. A Fluorescence in Situ Hybridization Method To Quantify mRNA Translation by Visualizing Ribosome–mRNA Interactions in Single Cells. ACS Cent. Sci. 3, 425–433 (2017).

13 Wadsworth, G. M., Parikh, R. Y., Kim, H. D. & Choy, J. S. mRNA detection in budding yeast with single fluorophores. Nucleic Acids Res. 45, e141–e141 (2017).

14 Gaspar, I. et al. Quantitative mRNA Imaging with Dual Channel qFIT Probes to Monitor Distribution and Degree of Hybridization. ACS Chem. Biol. 13, 742–749 (2018).

15 He, L. et al. Fluorescence Resonance Energy Transfer-Based DNA Tetrahedron Nanotweezer for Highly Reliable Detection of Tumor-Related mRNA in Living Cells. ACS Nano 11, 4060–4066 (2017).

16 Wang, B., Chen, Z., Ren, D. & You, Z. A novel dual energy transfer probe for intracellular mRNA detection with high robustness and specificity. Sensor. Actuat. B-Chem. 279, 342–350 (2019).

17 Chen, X. et al. Visualizing RNA dynamics in live cells with bright and stable fluorescent RNAs. Nat. Biotechnol. 37, 1287–1293 (2019).

18 Urbanek, M. O., Galka-Marciniak, P., Olejniczak, M. & Krzyzosiak, W. J. RNA imaging in living cells – methods and applications. RNA Biol 11, 1083–1095 (2014).

19 Bertrand, E. et al. Localization of ASH1 mRNA Particles in Living Yeast. Molecular Cell 2, 437–445 (1998).

20 Yan, X., Hoek, T. A., Vale, R. D. & Tanenbaum, M. E. Dynamics of Translation of Single mRNA Molecules In Vivo. Cell 165, 976–989 (2016).

21 Wang, C., Han, B., Zhou, R. & Zhuang, X. Real-Time Imaging of Translation on Single mRNA Transcripts in Live Cells. Cell 165, 990–1001 (2016).

22 Mamot, A. et al. Azido-Functionalized 5′ Cap Analogues for the Preparation of Translationally Active mRNAs Suitable for Fluorescent Labeling in Living Cells. Angew. Chem. Int. Ed. 56, 15628–15632 (2017).

23 Anhäuser, L., Hüwel, S., Zobel, T. & Rentmeister, A. Multiple covalent fluorescence labeling of eukaryotic mRNA at the poly(A) tail enhances translation and can be performed in living cells. Nucleic Acids Res. 47, e42–e42 (2019).

24 Croce, S., Serdjukow, S., Carell, T. & Frischmuth, T. Chemoenzymatic Preparation of Functional Click-Labeled Messenger RNA. ChemBioChem 21, 1641–1646 (2020).

25 Armitage, B. A. in Heterocyclic Polymethine Dyes: Synthesis, Properties and Applications 11–29 (Springer Berlin Heidelberg, Berlin, 2008).

26 Zhang, Y. & Kleiner, R. E. A Metabolic Engineering Approach to Incorporate Modified Pyrimidine Nucleosides into Cellular RNA. J. Am. Chem. Soc. 141, 3347–3351 (2019).

27 Ziemniak, M. et al. Synthesis and evaluation of fluorescent cap analogues for mRNA labelling. RSC advances 3, 20943–20958 (2013).

28 Xu, W., Chan, K. M. & Kool, E. T. Fluorescent nucleobases as tools for studying DNA and RNA. Nat. Chem. 9, 1043 (2017).

29 Sinkeldam, R. W., Greco, N. J. & Tor, Y. Fluorescent Analogs of Biomolecular Building Blocks: Design, Properties, and Applications. Chem. Rev. 110, 2579–2619 (2010).

30 Wilhelmsson, L. M. Fluorescent nucleic acid base analogues. Q. Rev. Biophys. 43, 159–183 (2010).

31 Börjesson, K. et al. Nucleic Acid Base Analog FRET-Pair Facilitating Detailed Structural Measurements in Nucleic Acid Containing Systems. J. Am. Chem. Soc. 131, 4288–4293 (2009).

32 Füchtbauer, A. F. et al. Fluorescent RNA cytosine analogue – an internal probe for detailed structure and dynamics investigations. Sci. Rep. 7, 2393 (2017).

33 Füchtbauer, A. F. et al. Interbase FRET in RNA: from A to Z. Nucleic Acids Res. 47, 9990–9997 (2019).

34 Liu, W. et al. Stringent Nucleotide Recognition by the Ribosome at the Middle Codon Position. Molecules (Basel, Switzerland) 22, 1427 (2017).

35 Stengel, G., Urban, M., Purse, B. W. & Kuchta, R. D. Incorporation of the Fluorescent Ribonucleotide Analogue tCTP by T7 RNA Polymerase. Anal. Chem. 82, 1082–1089 (2010).

36 McCoy, L. S., Shin, D. & Tor, Y. Isomorphic Emissive GTP Surrogate Facilitates Initiation and Elongation of in Vitro Transcription Reactions. J. Am. Chem. Soc. 136, 15176–15184 (2014).

37 Li, Y., Fin, A., McCoy, L. & Tor, Y. Polymerase-Mediated Site-Specific Incorporation of a Synthetic Fluorescent Isomorphic G Surrogate into RNA. Angew. Chem. Int. Ed. 56, 1303–1307 (2017).

38 Tanpure, A. A. & Srivatsan, S. G. A Microenvironment-Sensitive Fluorescent Pyrimidine Ribonucleoside Analogue: Synthesis, Enzymatic Incorporation, and Fluorescence Detection of a DNA Abasic Site. Chem. Eur. J. 17, 12820–12827 (2011).

39 Manna, S. & Srivatsan, S. G. Synthesis and Enzymatic Incorporation of a Responsive Ribonucleoside Probe That Enables Quantitative Detection of Metallo-Base Pairs. Org. Lett. 21, 4646–4650 (2019).

40 Yoshikawa, M., Kato, T. & Takenishi, T. Studies of Phosphorylation. III. Selective Phosphorylation of Unprotected Nucleosides. Bulletin of the Chemical Society of Japan 42, 3505–3508 (1969).

41 Ludwig, J. & Eckstein, F. Rapid and efficient synthesis of nucleoside 5’-0-(1-thiotriphosphates), 5’-triphosphates and 2’,3’-cyclophosphorothioates using 2-chloro-4H-1,3,2-benzodioxaphosphorin-4-one. J. Org. Chem. 54, 631–635 (1989).

42 Roy, B., Depaix, A., Périgaud, C. & Peyrottes, S. Recent Trends in Nucleotide Synthesis. Chem. Rev. 116, 7854–7897 (2016).

43 Burgess, K. & Cook, D. Syntheses of Nucleoside Triphosphates. Chem. Rev. 100, 2047–2060 (2000).

44 Flamme, M., McKenzie, L. K., Sarac, I. & Hollenstein, M. Chemical methods for the modification of RNA. Methods 161, 64–82 (2019).

45 Hocek, M. Enzymatic Synthesis of Base-Functionalized Nucleic Acids for Sensing, Cross-linking, and Modulation of Protein–DNA Binding and Transcription. Accounts Chem. Res. 52, 1730–1737 (2019).

46 Gaur, R. K., Sproat, B. S. & Krupp, G. Novel solid phase synthesis of 2’-o-methylribonucleoside 5’-triphosphates and their α-thio analogues. Tet. Lett. 33, 3301–3304 (1992).

47 Schoetzau, T., Holletz, T. & Cech, D. A facile solid phase synthesis of 2′- and 3′-aminonucleoside triphosphates. Chem. Comm., 387–388 (1996).

48 Sarac, I. & Meier, C. Efficient Automated Solid-Phase Synthesis of DNA and RNA 5′-Triphosphates. Chem. Eur. J. 21, 16421–16426 (2015).

49 Warnecke, S. & Meier, C. Synthesis of Nucleoside Di- and Triphosphates and Dinucleoside Polyphosphates with cycloSal-Nucleotides. J. Org. Chem. 74, 3024–3030 (2009).

50 Tonn, V. C. & Meier, C. Solid-Phase Synthesis of (Poly)phosphorylated Nucleosides and Conjugates. Chem. Eur. J. 17, 9832–9842 (2011).

51 Sarac, I. & Hollenstein, M. Terminal Deoxynucleotidyl Transferase in the Synthesis and Modification of Nucleic Acids. ChemBioChem 20, 860–871 (2019).

52 Motea, E. A. & Berdis, A. J. Terminal deoxynucleotidyl transferase: The story of a misguided DNA polymerase. Biochimica et Biophysica Acta (BBA) – Proteins and Proteomics 1804, 1151–1166 (2010).

53 Thomas, C. et al. One-step enzymatic modification of RNA 3’ termini using polymerase θ. Nucleic Acids Res. 47, 3272–3283 (2019).

54 Winz, M.-L., Samanta, A., Benzinger, D. & Jäschke, A. Site-specific terminal and internal labeling of RNA by poly(A) polymerase tailing and copper-catalyzed or copper-free strain-promoted click chemistry. Nucleic Acids Res. 40, e78–e78 (2012).

55 Srivatsan, S. G. & Tor, Y. Fluorescent Pyrimidine Ribonucleotide: Synthesis, Enzymatic Incorporation, and Utilization. J. Am. Chem. Soc. 129, 2044–2053 (2007).

56 Vaught, J. D., Dewey, T. & Eaton, B. E. T7 RNA Polymerase Transcription with 5-Position Modified UTP Derivatives. J. Am. Chem. Soc. 126, 11231–11237 (2004).

57 Aurup, H. et al. Translation of 2’-modified mRNA in vitro and in vivo. Nucleic Acids Res. 22, 4963–4968 (1994).

58 Smith, C. C., Hollenstein, M. & Leumann, C. J. The synthesis and application of a diazirine-modified uridine analogue for investigating RNA–protein interactions. RSC Advances 4, 48228–48235 (2014).

59 Domnick, C., Eggert, F. & Kath-Schorr, S. Site-specific enzymatic introduction of a norbornene modified unnatural base into RNA and application in post-transcriptional labeling. Chem. Comm. 51, 8253–8256 (2015).

60 Temiakov, D. et al. Structural Basis for Substrate Selection by T7 RNA Polymerase. Cell 116, 381–391 (2004).

61 Lakowicz, J. R. in Principles of Fluorescence Spectroscopy (Springer, Boston, 2006).

62 Stengel, G., Urban, M., Purse, B. W. & Kuchta, R. D. High Density Labeling of Polymerase Chain Reaction Products with the Fluorescent Base Analogue tC^O^. Anal. Chem. 81, 9079–9085 (2009).

63 Stengel, G. et al. Ambivalent Incorporation of the Fluorescent Cytosine Analogues tC and tC^O^ by Human DNA Polymerase α and Klenow Fragment. Biochemistry 48, 7547–7555 (2009).

64 Noyon, C. et al. The presence of modified nucleosides in extracellular fluids leads to the specific incorporation of 5-chlorocytidine into RNA and modulates the transcription and translation. Mol. Cell. Biochem. 429, 59–71 (2017).

65 Wittrup, A. et al. Visualizing lipid-formulated siRNA release from endosomes and target gene knockdown. Nat. Biotechnol. 33, 870 (2015).

66 Martín-Montañez, E. et al. Efficiency of gene transfection reagents in NG108-15, SH-SY5Y and CHO-K1 cell lines. Methods Find. Exp. Clin. Pharmacol. 32, 291–297 (2010).

67 Ducho, C. et al. Bis-cycloSal-d4T-monophosphates: Drugs That Deliver Two Molecules of Bioactive Nucleotides. J. Med. Chem. 50, 1335–1346 (2007).

68 Eaton, D. F. in Pure and Applied Chemistry Vol. 60 1107 (1988).

69 Sousa, R. in Methods in Enzymology 65–74 (Academic Press, 2000).

70 Trevino, S. G. et al. Evolution of functional nucleic acids in the presence of nonheritable backbone heterogeneity. Proceedings of the National Academy of Sciences 108, 13492 (2011).

71 Meyer, A. J. et al. Transcription yield of fully 2′-modified RNA can be increased by the addition of thermostabilizing mutations to T7 RNA polymerase mutants. Nucleic Acids Res. 43, 7480–7488 (2015).

72 Binning, J. M. et al. Development of RNA Aptamers Targeting Ebola Virus VP35. Biochemistry 52, 8406–8419 (2013).

73 Crooks, G. E., Hon, G., Chandonia, J.-M. & Brenner, S. E. WebLogo: a sequence logo generator. Genome Res. 14, 1188–1190 (2004).

74 Thomen, P. et al. T7 RNA polymerase studied by force measurements varying cofactor concentration. Biophys. J. 95, 2423–2433 (2008).

75 Nicosia, A., Tagliavia, M. & Costa, S. Regeneration of total RNA purification silica-based columns. Biomed. Chromatogr. 24, 1263–1264 (2010).

76 Aranda, P. S., LaJoie, D. M. & Jorcyk, C. L. Bleach gel: a simple agarose gel for analyzing RNA quality. Electrophoresis 33, 366–369 (2012).

77 R: A language and environment for statistical computing. (R Core Team, 2013).

78 Schindelin, J. et al. Fiji: an open-source platform for biological-image analysis. Nat. Methods 9, 676–682 (2012).

79 Lawrence, A.-M. & Besir, H. U. S. Staining of proteins in gels with Coomassie G-250 without organic solvent and acetic acid. J. Vis. Exp. 30, 1350 (2009).

