## Supplementary Information for "Stealth fluorescence labeling for live microscopy imaging of mRNA delivery"

**Supplementary Table 1**. S2

**Supplementary Figure 1.** S3

**Supplementary Figure 2**. S4

**Supplementary Figure 3**. S5

**Supplementary Figure 4**. S6

**Supplementary Figure 5**. S7

**Supplementary Figure 6**. S8

**Supplementary Figure 7.** S9

**Supplementary Figure 8**. S10

**Supplementary Table 2.** S11

| Name | Sequence (5' to 3')[a] |
|---|---|
| **T2** | TAA TAC GAC TCA CTA TAG |
| **B2** | ATT ATG CTG AGT GAT ATC CTG TAC TCC TAA TGG GTA CA |
| **DNA 3** | ATT ATG CTG AGT GAT ATC CTT CTC CTT CAC TCC TCT C |
| **DNA 4** | ATT ATG CTG AGT GAT ATC CTA CTC CTT CCT TCC TCT C |
| **T12.3** | GCC TGT TGT GAG CCT CCT AAC GCA CCG GTC GCA GGT TTA GCA AGG TCA CAA GCT GGC ATC AAG GTT GCC CCA TGC TTA TTC TTG TCT CCC |
| **D1 Library** | GCC TGT TGT GAG CCT CCT AAC $N_{49}$ CA TGC TTA TTC TTG TCT CCC |
| **FWD-D1-RNA 2** | TAA TAC GAC TCA CTA TAG GGA GAC AAG AAT AAG CAT G |
| **REV-D1-RNA 2** | GCC TGT TGT GAG CCT CCT AAC |
| **FWD-D1-RNA 1** | TAA TAC GAC TCA CTA TAG GCC TGT TGT GAG CCT CCT AAC |
| **REV-D1-RNA 1** | GGG AGA CAA GAA TAA GCA TG |
| **REV-D1 128** | GCC TGT TGT GAG CCT CCT AAC |
| **REV_D1 23** | GGG AGA CAA GAA TAA GCA TG |
| **M13 uni (-43)** | AGG GTT TTC CCA GTC ACG ACG TT |
| **M13 rev (-29)** | CAG GAA ACA GCT ATG ACC |
| **TdT1** | FAM-GCA AGC ACA GAC AUC AG |
| **TdT2** | FAM-GGG AAG UGC UAC CAC AAC UUU AGC CAU AAU GUC ACU UCU GCC GCG GGC AU |
| **TdT3[b]** | GGG AAG UGC UAC CAC AAC UUU AGC CAU AAU GUC ACU UCU GCC GCG GGC AUG CGG CCA GCC A |

[a] the T7 promoter sequence is underlined; [b] obtained by transcription reaction.

**Supplementary Table 1.** Oligonucleotide sequences.

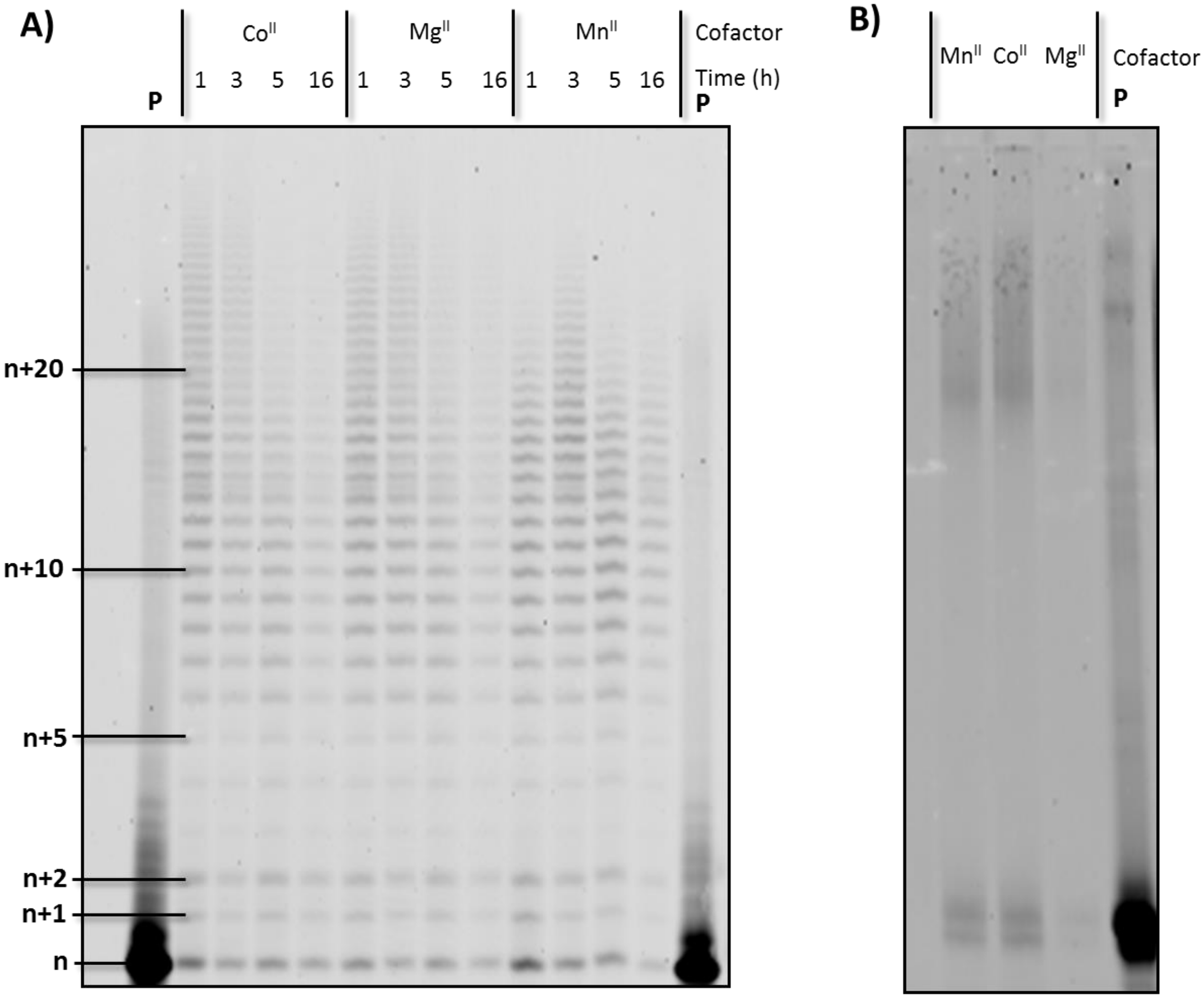


**Supplementary Figure 1**. Gel images (PAGE 20%) of the products stemming from the TdT-mediated tailing reactions with the modified triphosphate carried out using A) primer **TdT1** (also shown in the manuscript) and B) **TdT2**, different metal cofactors and reaction times. P indicates the unreacted primer.

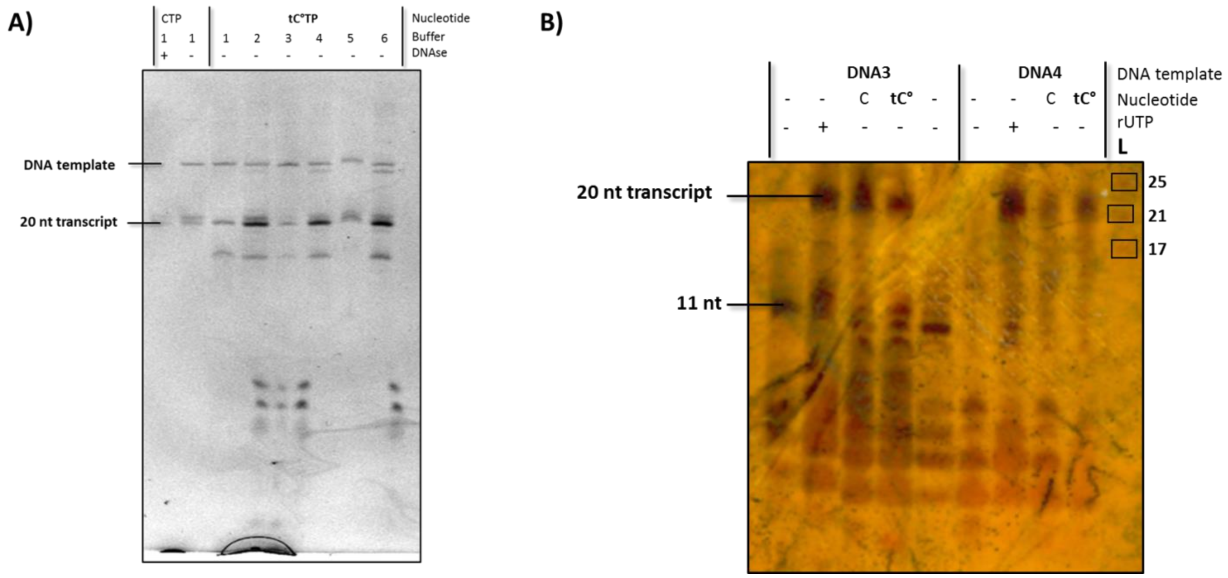


**Supplementary Figure 2**. Gel images (PAGE 20%) of transcription reactions with CTP and tC°TP. A) reactions performed with DNA oligonucleotides B2 and T2 and different buffer conditions (see Methods for buffer compositions). Midori green was used for visualization. B) transcription reactions with DNA templates DNA3 and DNA4. Control reactions were performed in the presence or absence of rUTP and all reactions were carried out in buffer 5 at 37°C for 30 min. L represents a ladder and visualization was performed by silver staining.

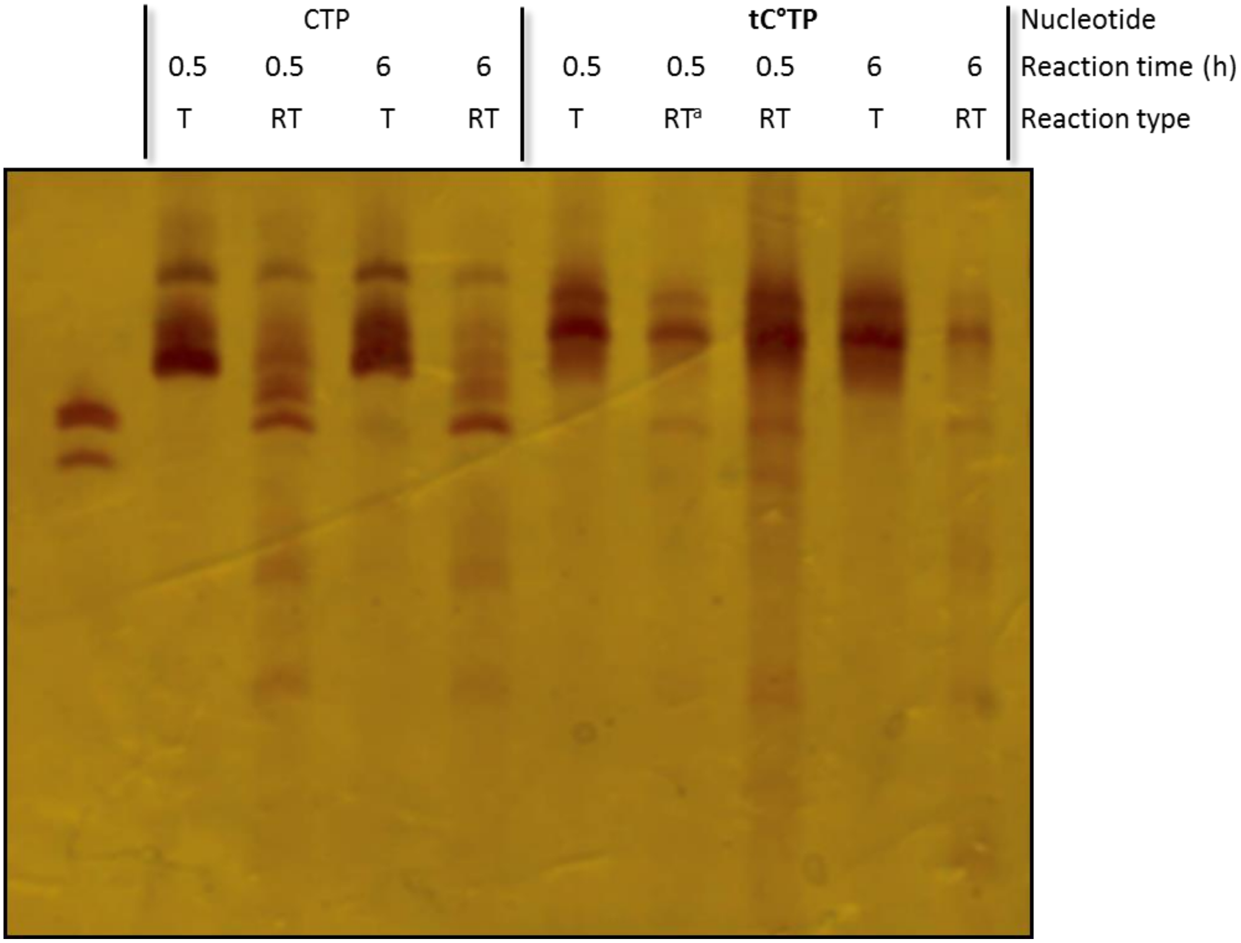


**Supplementary Figure 3**. Gel analysis (PAGE 10%) of the products stemming from transcription (T) from the T12.3 DNA template and subsequent reverse transcription (RT) reactions. The reaction times (0.5 and 6 h) refer to the times used for the transcription reactions. The leftmost lane shows the initial T12.3 DNA template. Visualization by silver staining. [a]5x less material was loaded on the gel.

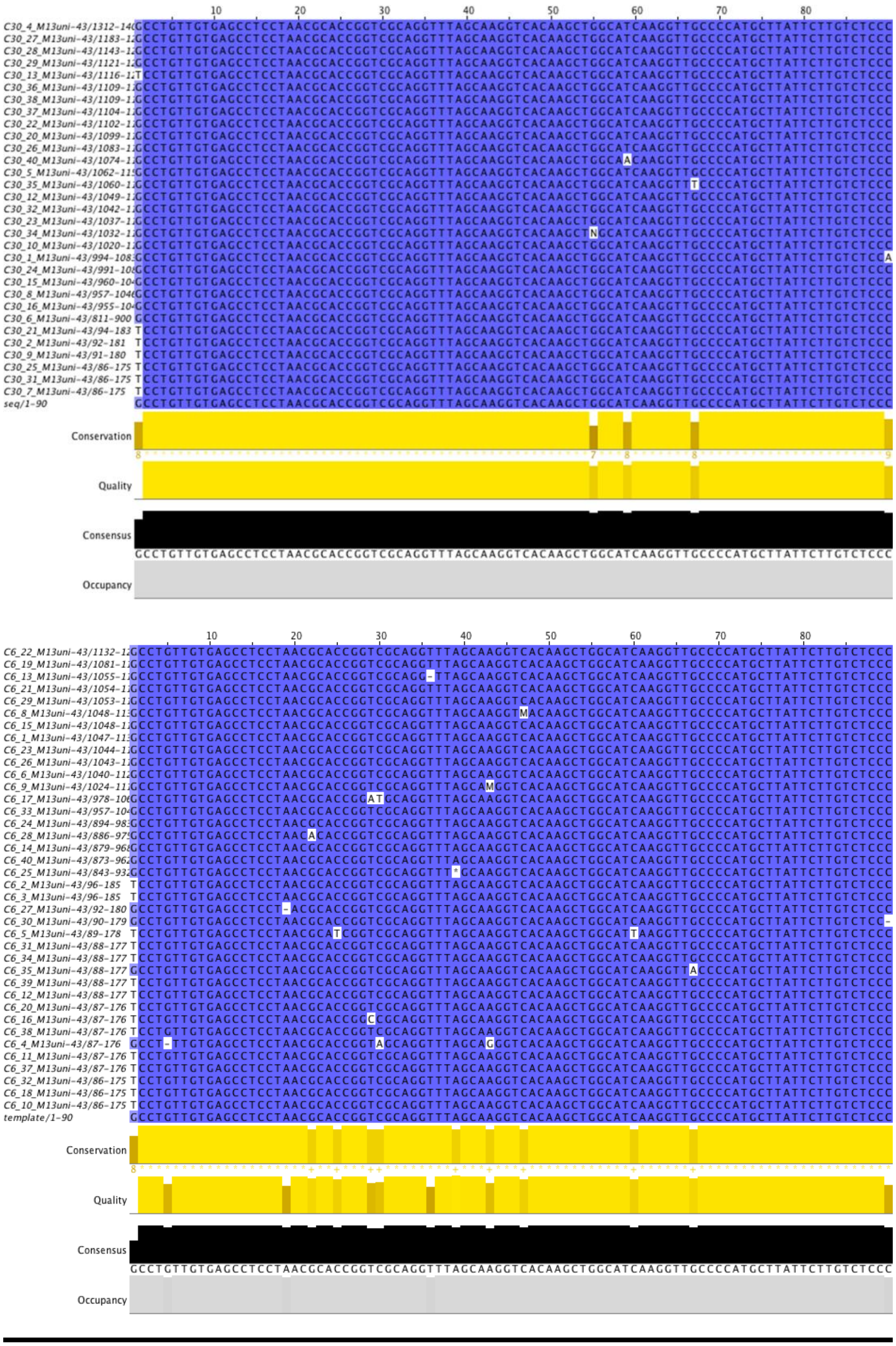


**Supplementary Figure 4**. Multiple alignment of the sequences stemming from the transcription reactions with CTP (up: 30 min and down: 6 h reaction time). Y: C or T, S: G or C, R: A or G, W: A or T, K: G or T, M: A or C, N: any nucleotide.

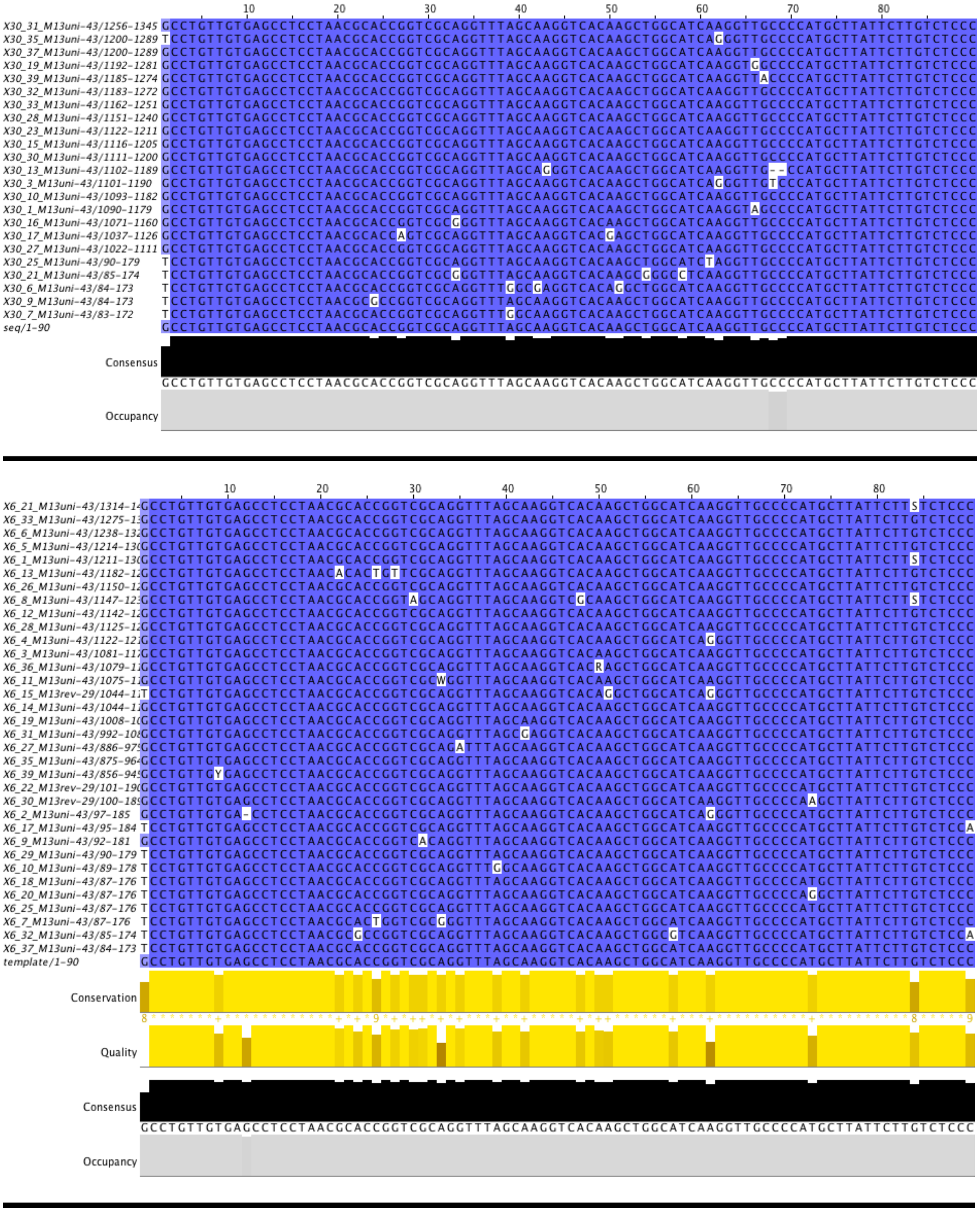

**Supplementary Figure 5**. Multiple alignment of the sequences stemming from the transcription reactions with tC°TP (up: 30 min and down: 6 h reaction time). Y: C or T, S: G or C, R: A or G, W: A or T, K: G or T, M: A or C, N: any nucleotide.

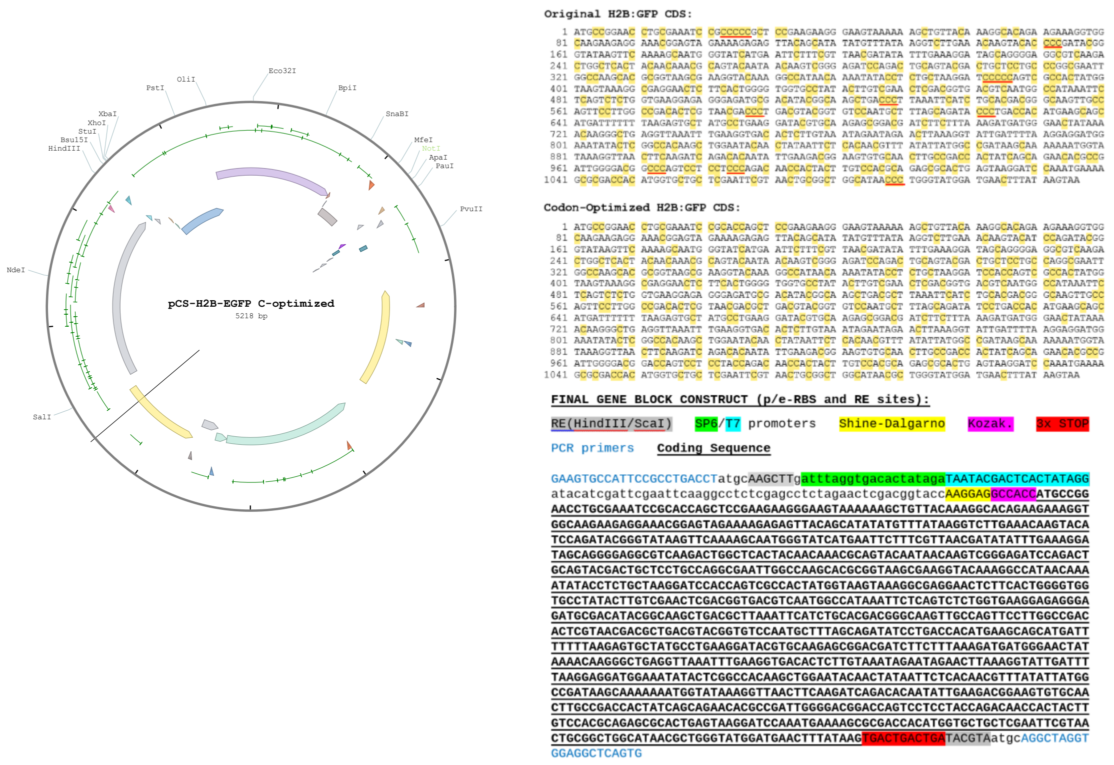


**Supplementary Figure 6**. pCS2-H2B:GFP plasmid, optimization of the H2B:GFP template and final construct.

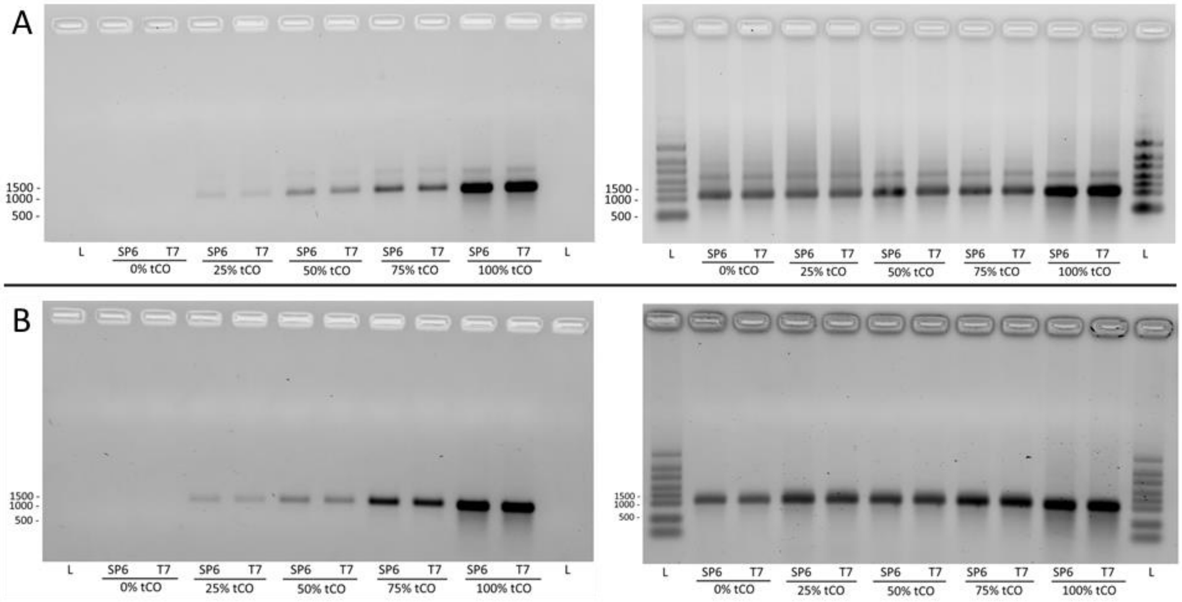


**Suplementary Figure 7.** Incorporation of tC$^{O}$ into full length mRNA by SP6 and T7 RNA polymerase assisted in vitro transcription. (a) Denaturing agarose bleach gels showing RNA transcripts formed at five different tC$^{O}$TP/CTP ratios (0-100%). The produced RNA was visualized directly by tC$^{O}$ fluorescence (left image) and after ethidium bromide staining (right image). The RNA samples were heat-denatured (65 °C for 5 min, 1.5 % bleach in the gel) prior to loading on the gel. (b) Denaturing bleach gels of the same RNA transcripts as in (A) but at stronger denaturing conditions (70 °C for 10 min., 2 % bleach in the gel). The RiboRuler High Range RNA ladder was used.

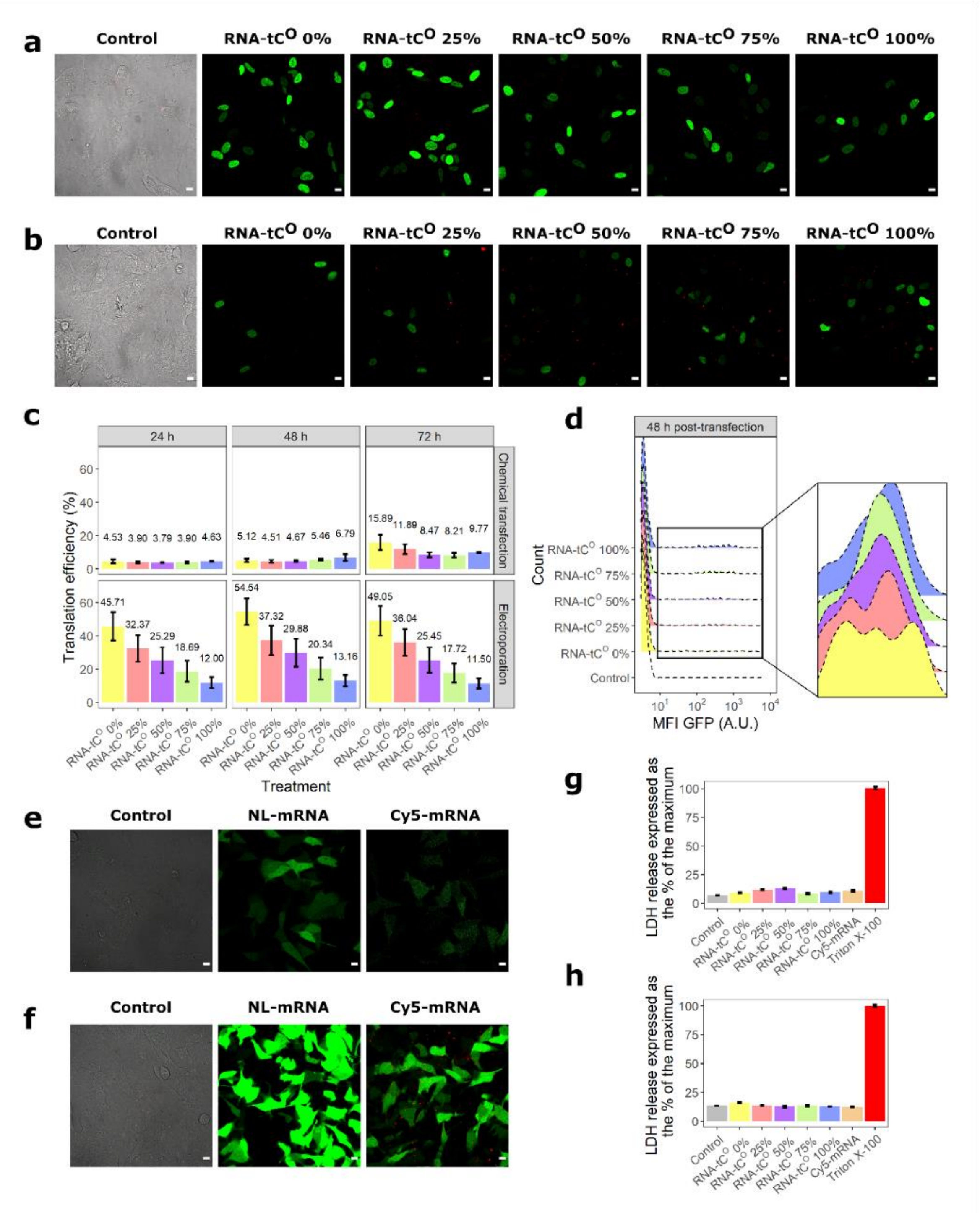


**Supplementary Figure 8. Translation efficiency of the modified RNA constructs in human cells and cytotoxicity assessment.** Representative confocal images (large view, scale bar: 10 µm) of RNA-tC$^{O}$ constructs and mRNAs from Trillink® transfected by (a, e) electroporation or (b, f) chemical transfection. (c) Percentages of positive cells for H2B:GFP at 24 h, 48 h and 72 h post-transfection with RNA-tC$^{O}$ constructs. (d) Representative histogram of the GFP signal distribution in single living cells at 48 h post-chemical transfection. Cytotoxicity assessment performed 24 h (g) post-electroporation or (h) post-chemical transfection using the LDH cell membrane integrity assay.

| | transcript | $A_1$ | $T_1$ (ns) | $A_2$ | $T_2$ (ns) | $A_3$ | $T_3$ (ns) | $\bar{\tau}$ (ns) | $X^2$ |
|---|---|---|---|---|---|---|---|---|---|
| SET 1 | 25% | 0.19 | 0.67 | 0.49 | 3.2 | 0.31 | 5.6 | 3.5 | 1.09 |
| | 50% | 0.28 | 0.71 | 0.43 | 2.8 | 0.30 | 5.2 | 2.9 | 1.06 |
| | 75% | 0.33 | 0.68 | 0.45 | 2.6 | 0.23 | 5.2 | 2.5 | 1.12 |
| | 100% | 0.33 | 0.48 | 0.44 | 2.1 | 0.23 | 4.8 | 2.2 | 1.00 |
| SET 2 | 25% | 0.21 | 0.63 | 0.40 | 2.8 | 0.34 | 5.3 | 3.3 | 0.99 |
| | 50% | 0.26 | 0.64 | 0.41 | 2.7 | 0.30 | 5.2 | 3.0 | 1.03 |
| | 75% | 0.29 | 0.50 | 0.42 | 2.1 | 0.22 | 4.8 | 2.4 | 1.02 |
| | 100% | 0.37 | 0.47 | 0.40 | 1.9 | 0.39 | 4.5 | 2.0 | 1.01 |

**Supplementary Table 2**. Fitted lifetime parameters for the TCSPC experiments. The $X^2$-value (Chi-Square) was evaluated to indicate goodness of fit.
